# A novel, high-density CRISPR activation platform for mapping cancer dependencies and resistance pathways *ex vivo* and *in vivo*

**DOI:** 10.1101/2025.07.22.666069

**Authors:** Sarah T. Diepstraten, Yexuan Deng, Margaret A. Potts, Amy Heidersbach, Christina König, Kristel M. Dorighi, Lin Tai, Andrew J. Kueh, Lauren Whelan, Catherine Chang, Felix Brown, Gemma L. Kelly, Jean-Philippe Fortin, Benjamin Haley, John E. La Marca, Marco J. Herold

**Author notes:** corresponding author: Marco Herold. Université de Montréal, Centre de recherche de l’Hôpital Maisonneuve-Rosemont (Benjamin Haley). these authors share first author status (Sarah T. Diepstraten, Yexuan Deng, Margaret A. Potts, Amy Heidersbach). these authors share senior author status (Gemma L. Kelly, Jean-Philippe Fortin, Benjamin Haley, John E. La Marca, Marco J. Herold).

## Abstract

CRISPR activation (CRISPRa) enables precise, locus-specific upregulation of gene expression, offering potential for both *ex vivo* and *in vivo* applications. However, the lack of scalable, high-coverage tools has limited its use in comprehensive genetic screens, particularly in murine models. Here, we introduce Partita, a next-generation, whole-genome CRISPRa sgRNA platform designed for unparalleled efficiency and depth in gene activation studies. Partita employs a high-density targeting strategy, deploying 10 sgRNAs per transcription start site, structured into five gene family-specific sub-libraries to maximize transcriptional induction. To demonstrate its capabilities, we performed a series of large-scale screens: an *in vitro* enrichment/depletion screen in iBMDMs, whole-genome CRISPRa screens in a double-hit lymphoma model to uncover genes driving resistance to pro-apoptotic drugs (venetoclax, nutlin-3a, etoposide), and an *in vivo* whole-genome screen identifying accelerators of Myc-driven lymphomagenesis. Each experiment revealed both expected and novel regulators of cellular phenotypes, with a high validation rate in secondary assays. By enabling robust, high-throughput gain-of-function screening, Partita unlocks new avenues for functional genomics and expands the toolkit for discovering key drivers of biological processes across diverse research fields.

## Introduction

Clustered regularly interspaced short palindromic repeat activation (CRISPRa) is an adaptation of CRISPR technology that enables target-specific gene expression. Canonical CRISPR technology uses a Cas nuclease protein (e.g. Cas9) to generate DNA double-stranded breaks, as directed by a single guide RNA (sgRNA) to the locus of interest (*1*). CRISPRa differs as it employs a nuclease-dead Cas protein (e.g. dCas9) engineered with or capable of recruiting transcriptional activation domains (TADs) (*2*). When coupled with sgRNAs that direct the nuclease-dead Cas protein to the “promoter” regions upstream of the transcriptional start site (TSS), gene induction is enabled. A variety of CRISPRa modalities have been developed, such as dCas9-SAM (*3*), dCas9-VPR (*4*), or SunTag (*5*), which differ in aspects like the TADs used and sgRNA scaffold designs (*2*). A number of mouse models for CRISPRa have also been generated, including some using the SunTag (*6, 7*) and dCas9-Synergistic Activation Mediator (SAM) systems (*8–10*). These models have greatly increased the utility of CRISPRa technology *in vitro* and *in vivo*, as they permit the bypassing of complicated construct delivery into primary mouse cells.

One of the strengths of CRISPR technology has been the ability to conduct large-scale genetic screens with relative ease. Several whole-genome CRISPRa sgRNA libraries are publicly available (*3, 11–13*), each with their own advantages and disadvantages. CRISPRa screening is less straightforward than traditional CRISPR knock-out screening, as precise targeting of transcriptional start sites (TSS) is necessary to achieve gene activation. Accurately identifying gene TSS can be difficult, and is an ongoing process for many genes (*14*), making design of effective sgRNAs much more difficult than for CRISPR knock-out. In addition, unlike CRISPR knock-out which requires only a single instance of gene editing to induce a permanent mutation, CRISPRa requires constitutive and high levels of all elements of the CRISPRa system, including dCas9, TADs and sgRNAs, to initiate and maintain gene activation.

In this study, we have generated a new mouse CRISPRa whole-genome sgRNA platform – Partita – which is compatible with the dCas9-SAM system. To increase the probability of target gene activation, the library contains 10 sgRNAs per gene. Additionally, many previous methods of CRISPRa sgRNA library construction have exclusively directed the dCas9 activator to a single TSS per gene, which could potentially limit observations to phenotypes derived from isoform (TSS)- specific expression (*14, 15*). To account for this, we established an sgRNA targeting scheme using a new methodology informed by multiple datasets across various cell types of high-quality mouse TSS annotations, offering the potential for activating up to seven distinct promoters per gene. This results in a library with >260,000 sgRNAs covering >26,000 TSSs for 21,557 genes, with accompanying non-targeting control (NTC) sgRNAs. To facilitate diverse screening applications, the Partita whole-genome library is separated into 5 sub-libraries: 1) DNA repair, epigenetics, GPCRs, kinases, phosphatases, proteases, ubiquitination; 2) ion channel, drug targets, secretome (part 1); 3) secretome (part 2); 4) others (part 1); and 5) others (part 2). We further generated a sixth sub-library, which collected only those sgRNAs targeting transcriptional modifiers from sub-libraries 1-5. Finally, the library backbone contains both puromycin resistance and BFP fluorescence selection markers for ease of use, along with a previously optimized sgRNA scaffold (*16*), to increase overall library functionality.

Herein, we validate this library in a number of biological contexts, including: conventional sgRNA enrichment/depletion screens in immortalised bone marrow-derived macrophages (iBMDMs), whole-genome screens to identify known and novel resistance factors to anti-cancer drugs in a mouse model of “double-hit” diffuse large B cell lymphoma, and whole-genome unbiased screening for and validation of oncogenes that co-operate with *Myc*-driven pre-B/B cell lymphoma development *in vivo*. We anticipate that these libraries will be broadly useful tools for the scientific community, enabling the study and discovery of new gene functions, disease biology, and therapies.

## Results

### Designing a next-generation CRISPRa sgRNA library

While sgRNA targeting schemes for CRISPRa have advanced in recent years, the ability to predict a function for each guide remains low compared to gene knock-out/suppression methods. Proper targeting of the transcriptional start site (TSS) is key to achieving robust gene upregulation (*13, 17–19*), and so we aimed to generate a murine CRISPRa whole genome sgRNA library using some of the most up-to-date TSS information. Mouse protein-coding genes were first identified using GENCODE Mouse Release 25 (GRCm38). Candidate TSSs were identified using the RefTSS database, which combines datasets including FANTOM5, EPDnew, and ENCODE (*20*), with a specific focus on identifying those TSSs located in a [-500 bp, 500 bp] window around the start of GENCODE-annotated isoforms. From these data, a primary promoter (P1; selected as the reference TSS and positioned closest to the 5ill end of the isoform annotated as “principal” by APPRIS) was nominated for each gene, as well as up to six additional promoters per gene (P2-P7), for a total of 24,522 promoter regions (P1 = 21,833; P2 = 2474; P3 = 197; P4 = 13; P5 = 3; P6 = 2; P7 = 1). Two additional sets of cell-type-specific TSSs were also included. Datasets from murine embryonic stem cells (mESCs) (GSE98063) (*21*) and murine bone marrow-derived macrophages (mBMDMs) (*22*) yielded 70 mESC-specific and 43 mBMDM-specific TSSs. Finally, we also included an additional 1557 promoters taken from the leading mouse CRISPRa whole-genome sgRNA libraries: Caprano (*13*) and Weissman v2 (*11*), provided that they were at least 1 kb removed from our already selected promoters. In total, we included 26,192 TSSs in our annotation.

Using this annotation as our foundation, we designed 10 sgRNAs per TSS. The sgRNAs with spacer sequences containing either EcoRI or KpnI restriction enzyme recognition sites were filtered out (to eliminate complications with library synthesis/cloning methods). As the distance between corresponding sgRNAs and TSSs has been shown to be a major determinant of guide efficacy (*13*), we prioritised the selection of sgRNAs located within a projected optimal activation window of [-150 bp, -75 bp], while also placing guides within an additional six regions of decreasing priority (and increasing size) to ensure 10 sgRNAs per promoter were obtained (Fig. S1A). Within each of these regions, sgRNAs were further filtered to remove those with a poly-T termination signal and extreme GC content, followed by ranking prospective sequences based on Azimuth on-target activity score. The collection of sgRNA sequences for each sub-library were then cloned as pools into the WEHI-12 vector (Fig. S1B), which contains a bi- cistronic BFP fluorescence and puromycin antibiotic resistance marker cassette for enhanced utility.

In total, 263,460 sgRNAs were included in the final whole-genome library, collectively named “Partita”. The substantial volume of sgRNAs comprising the full Partita library could be restrictive for some screening applications. Therefore, we created five non-overlapping sub-libraries (each with 1000 NTC sgRNAs): sub-library 1 contains sgRNAs targeting genes involved in DNA repair, ubiquitination, and epigenetic mechanisms, G protein-coupled receptor genes, kinase genes, phosphatase genes, and protease genes; sub-library 2 contains ion channel-related genes and genes implicated as encoding druggable targets; sub-library 3 contains genes encoding the secretome, and sub-libraries 4 and 5 contain all other remaining genes (Fig. 1, Table 1). Additionally, we generated a sixth boutique Partita sub-library, which combined all sgRNAs from sub-libraries 1-5 that target transcription modifiers (Fig. S1A). Our sub-library concept enables both unbiased genome-wide screening using all libraries, or targeted screening of select genes/gene families with maximized TSS coverage. To demonstrate the efficacy and broad utility of the Partita library, we then employed it in a range of screening experiments *in vitro* and *in vivo*.

**Figure 1.**
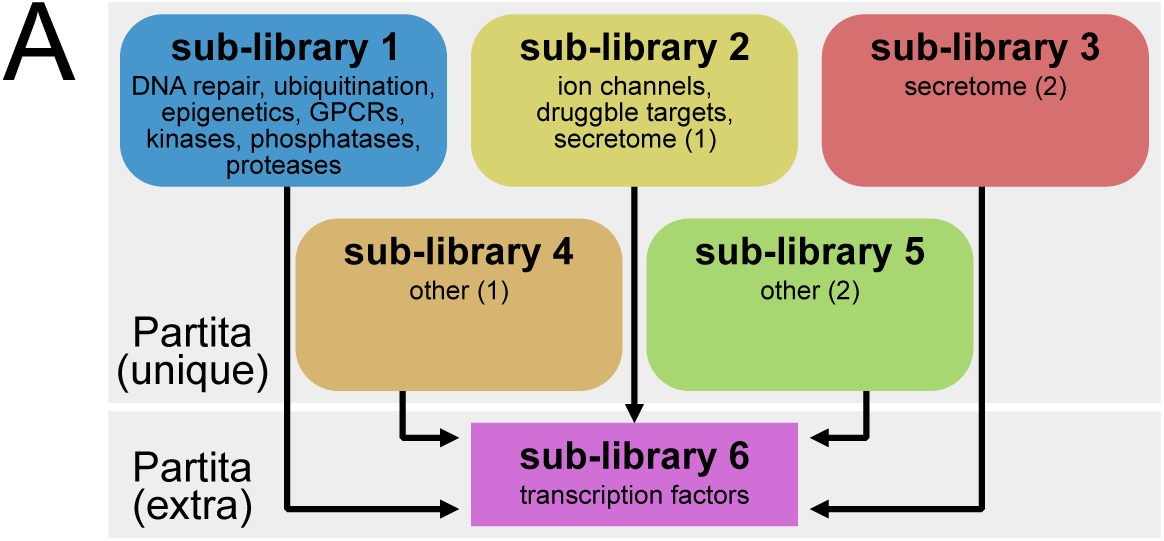
Partita whole-genome CRISPRa sgRNA library overview. (A) The Partita whole-genome library is separated into 5 sub-libraries. Sub-library 1 contains sgRNAs targeting genes related to DNA repair, epigenetics, GPCRs, kinases, phosphatases, proteases, and ubiquitination. Sub-library 2 contains sgRNAs targeting genes related to ion channels, drug targets, and the secretome (part 1); Sub-library 3 contains sgRNAs targeting genes related to the secretome (part 2). Sub-libraries 4 and 5 contain all other genes (parts 1 and 2). Additionally, a sixth sub-library contains only those sgRNAs targeting transcriptional modifiers, that are taken from sub-libraries 1-5 (therefore, this sub-library is not unique). Each library also contains 1000 NTC sgRNAs.

**Table 1.** Partita sub-library composition. Details of the gene classes targeted by the sgRNAs in each sub-library of the Partita whole-genome sgRNA library, as well as the number of unique genes targeted by each sub-library and the number of unique sgRNAs. Note that sub-library 6, which targets only transcriptional modifier genes, uses sgRNAs taken from the other sub-libraries. Complete library information can be found in Supplementary File 1.

| Partita sub-library | Sub-library gene targets | Number of target genes | Number of sgRNAs |
| --- | --- | --- | --- |
| 1 | DNA repair<br>Ubiquitination<br>Epigenetics<br>G protein-coupled receptors<br>Kinases<br>Phosphatases<br>Proteases | 4506 | 53,650<br>(+1000 NTC) |
| 2 | Ion channels<br>Druggable targets<br>Secretome (1) | 4532 | 54,168<br>(+1000 NTC) |
| 3 | Secretome (2) | 4687 | 54,571<br>(+1000 NTC) |
| 4 | Other (1) | 4634 | 54,284<br>(+1000 NTC) |
| 5 | Other (2) | 3198 | 42,649<br>(+1000 NTC) |
| 6 | Transcriptional modifiers<br>(non-unique, extracted from other sub-libraries) | 2339 | 23,737<br>(+1000 NTC) |

### Cell growth modifier screen in immortalised bone marrow-derive macrophages

To benchmark the efficacy of the Partita libraries we performed a whole-genome enrichment/depletion CRISPRa screen, which aimed to identify genes that either promoted or inhibited cell growth upon activation (Fig. 2A). While general cell growth screens are common with CRISPR-based gene knock-out/suppression approaches, comparable CRISPRa screens have not yet been published in murine primary cells. This presented the opportunity for both discovery and validation of anticipated factors (e.g. *Trp53* as a negative growth regulator). To ensure consistent, stable expression of the CRISPRa machinery, we utilised cells derived from our previously characterised *dCas9a-SAM^KI/KI^* mouse model, in which cells constitutively express *dCas9-SAM* (*8*). From these mice, we generated immortalised bone marrow-derived macrophages (iBMDMs) using *Cre-J2* retrovirus, as described previously (*23*), creating a non-malignant cell line that is easy to expand and transduce (Fig. 2A). As we sought to perform the enrichment/depletion screen under normal conditions, without any additional cellular stress, we expected any top hits might be targeting genes that relate to cell proliferation (for enriched sgRNAs) or cell death/stasis (for depleted sgRNAs). Therefore, we chose to use Partita sub-library 1, which contains sgRNAs targeting several genes known to have functions related to cell survival and proliferation (Fig. 2A). To ensure our screen had sufficient statistical power to accurately assess the depletion of all sgRNAs, we aimed for 500x sgRNA coverage (i.e. 500 cells per sgRNA after library transduction (Figs. 2A, S2A)). Finally, we performed this screen across three biological replicates, in parallel, to further increase the reliability of the screen results (Fig. 2A). After transduction, the iBMDMs were treated with puromycin to select for successfully transduced cells, and post-puromycin selection the initial input sample was collected for DNA analysis (time point 0). The iBMDMs were then expanded and cultured under normal conditions for three weeks, with genomic DNA harvested from cell populations at the end of weeks one, two, and three (time points 1/2/3) (Fig. 2A). Each sample was then analysed by next generation sequencing (NGS) to quantify sgRNA levels and, by extension, gene-specific effects on cell growth.

**Figure 2.**
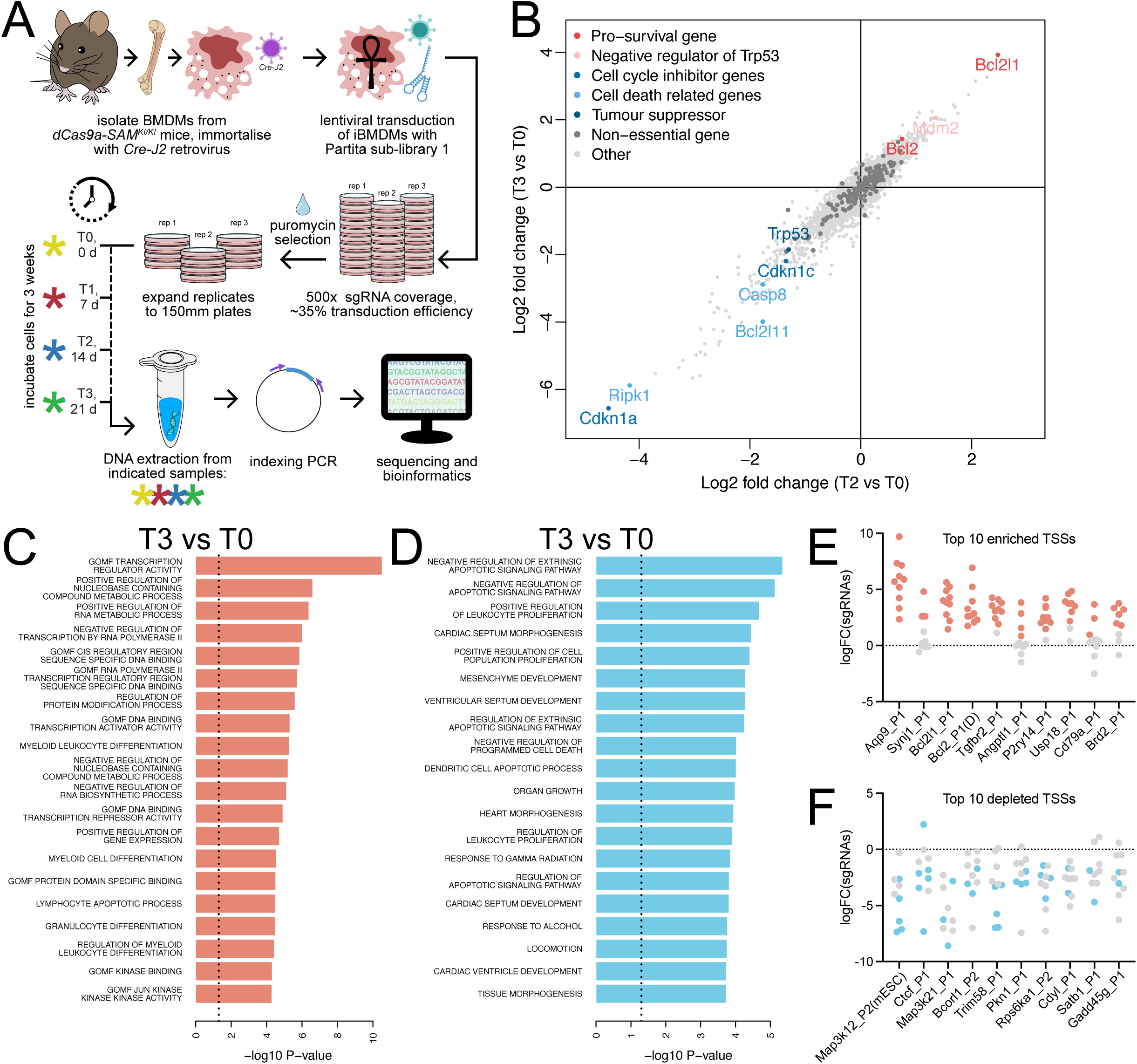
Enrichment/depletion screening in iBMDMs using Partita sub-library 1. (A) Diagram of the screening setup. BMDMs were first isolated from *dCas9a-SAM^KI/KI^* mice and immortalised using *Cre-J2* retroviral infection. iBMDMs were then lentivirally transduced with Partita sub-library 1, at a transduction efficiency of ∼35% in order to obtain 500x sgRNA coverage. Three replicate infections were performed, with 10 plates (100mm) per replicate. Puromycin was then used to select for successfully transduced cells, and the remaining cells from each replicate were then expanded into 4 larger plates (150mm). These plates were incubated for 3 weeks, with cells harvested at 4 time points (T0 = 0 days, T1 = 7 days, T2 = 14 days, T3 = 21 days). DNA was then extracted from each of these samples, before indexing PCR, next-generation sequencing, and bioinformatics analyses were performed. (B) Four-way plot showing gene enrichment and gene depletion, with T2 vs t0 on the x-axis, and T3 vs T0 on the y-axis. Pro-survival and anti-apoptotic genes are observed to be clearly enriched over time, while cell-cycle inhibitors and pro-apoptotic genes are observed to be depleted. (C) GSEA graphs showing the top twenty depleted gene classes at T3, relative to T0. Note the depletion of gene classes likely to be deleterious if activated, such as negative transcriptional regulators. Additional time points are shown in Fig.S2B. (D) GSEA graphs showing the top twenty enriched gene classes at T3, relative to T0. Note the enrichment of gene classes likely to be advantageous if activated, such as negative regulators of apoptosis. Additional time points are shown in Fig.S2C. (E) Plot showing the logFC of each sgRNA targeting each of the top 10 enriched TSSs at T3, relative to T0. Each dot represents an independent sgRNA. Significantly changed (FDR <0.05) sgRNAs are show in red. (F) Plot showing the logFC of each sgRNA targeting each of the top 10 depleted TSSs at T3, relative to T0. Each dot represents an independent sgRNA. Significantly changed (FDR <0.05) sgRNAs are shown in blue.

Out of 5306 TSSs potentially detectable at T0, when comparing T1 vs T0 we observed 382 enriched and 208 depleted TSSs, when comparing T2 vs T0 we observed 1017 enriched and 477 depleted TSSs, and when comparing T3 vs T0 we observed 1199 enriched and 522 depleted TSSs (thresholds of FDR<0.05 and logFC>0.25 for enriched TSSs, FDR<0.05 and logFC<-0.5 for depleted TSSs; count data available at GSE296642). A 4-way comparative analysis of sgRNA enrichment/depletion for T1 vs T0 and T3 vs T0 revealed a significant shift towards depletion in sgRNAs targeting genes known to promote cell death or cell senescence upon activation – such as *Trp53*, *Cdkn1a*, and *Casp8* – while non-targeting control sgRNAs remained relatively unchanged (Fig. 2B). Notably, for *Cdkn1a*, two individual sgRNAs (representing two separate TSS target loci) were observed as being significantly depleted. Consistent effects following activation of distinct promoters for this gene increased the confidence in our results and sgRNA targeting scheme. We also observed a shift towards enrichment of sgRNAs targeting genes that promote cell survival upon activation – such as *Bcl2l1* (encoding BCL-XL), *Bcl2*, and *Mdm2* (Fig. 2B), the latter of which encodes a potent negative regulator of TRP53 (*24–26*) (Fig. 2B). These observations are supported by gene set enrichment analyses (GSEAs), which showed that depleted gene classes were generally those that inhibit cell proliferation (Fig. 2C, S2B), whereas enriched gene classes were those that support cell survival (Fig. 2D, S2C). To examine sgRNA efficiency, we also looked at individual sgRNAs targeting the top 10 enriched and depleted TSSs in this screen. For each of the enriched TSSs, most of their sgRNAs were significantly enriched (Fig. 2E). For each of the depleted TSSs, fewer sgRNAs were significantly depleted, likely due to the increased difficulty of detecting depleted sgRNAs at a significant level, though multiple significantly depleted sgRNAs were still detected (Fig. 2F). Overall, these data indicate that the Partita sgRNAs are efficient and effective, and demonstrate the strength of the Partita libraries for use in a conventional CRISPR enrichment/depletion screening approach.

### Screening in double-hit lymphoma cells using the Partita library to identify resistance factors to venetoclax and other anti-cancer drugs

We next sought to employ the Partita library in a disease model, to identify novel mechanisms of therapeutic resistance or sensitisation. The transcription factor TRP53 (TP53 in humans) is a potent tumour suppressor that can drive induction of intrinsic apoptosis, cell cycle arrest, and cell senescence (*27*). The *TP53* gene is lost or mutated in 50% of human cancers (*28*). Identifying factors downstream of TRP53, which are necessary for induction of apoptosis, may reveal new targets to induce cell death in *TP53*-mutant cancers. To this end, we undertook a screen to identify novel regulators of TRP53-mediated apoptosis in a mouse model of aggressive “double-hit” lymphoma (DHL) model (*8*). Human double-hit lymphoma is a high-grade diffuse large B cell lymphoma defined by simultaneous upregulation of BCL2 and MYC (*29*). Our murine DHL model combines *Eµ-Myc* transgene expression (*30, 31*) with constitutive *dCas9-SAM* expression and a corresponding sgRNA to promote *Bcl2* upregulation. In our experience, DHL cell lines have proven difficult to transduce, making them a poor choice for depletion screening (which requires high coverage for appropriate statistical power), but an effective model for enrichment screening to identify genes promoting resistance to various anti-cancer therapies.

We first chose venetoclax (a.k.a. ABT-199), a BH3-mimetic that selectively inhibits BCL-2, the titular member of the BCL-2 family of pro-survival proteins (*32*). Venetoclax is FDA approved for the treatment of chronic lymphocytic leukaemia and acute myeloid leukaemia but, despite good upfront responses, many patients eventually relapse due to the development of secondary resistance (*33*). We have previously identified some resistance factors for venetoclax via CRISPRa screening in DHL cells using the Caprano library (*13*), such as compensatory upregulation of the untargeted pro-survival proteins MCL-1, BCL-XL, and BFL-1/A1 (*8*). Therefore, we sought to validate the use of our new Partita libraries in this context. Two DHL cell lines, DHL216 and DHL214, were transduced with Partita sub-libraries 1-5 individually. Flow cytometric analysis detected 10-15% successfully transduced, BFP-positive cells (Fig. S3A). These cells were then expanded for ∼10 days to enable maximal target gene induction. Cells were then treated with venetoclax (IC_50_ doses for 3 rounds of treatment, or IC_80_ doses for 1 round of treatment) or with DMSO as a vehicle control (Fig. 3A). The cells were then allowed to recover, before being collected and their DNA sequenced to quantify sgRNAs (Fig. 3A).

**Figure 3.**
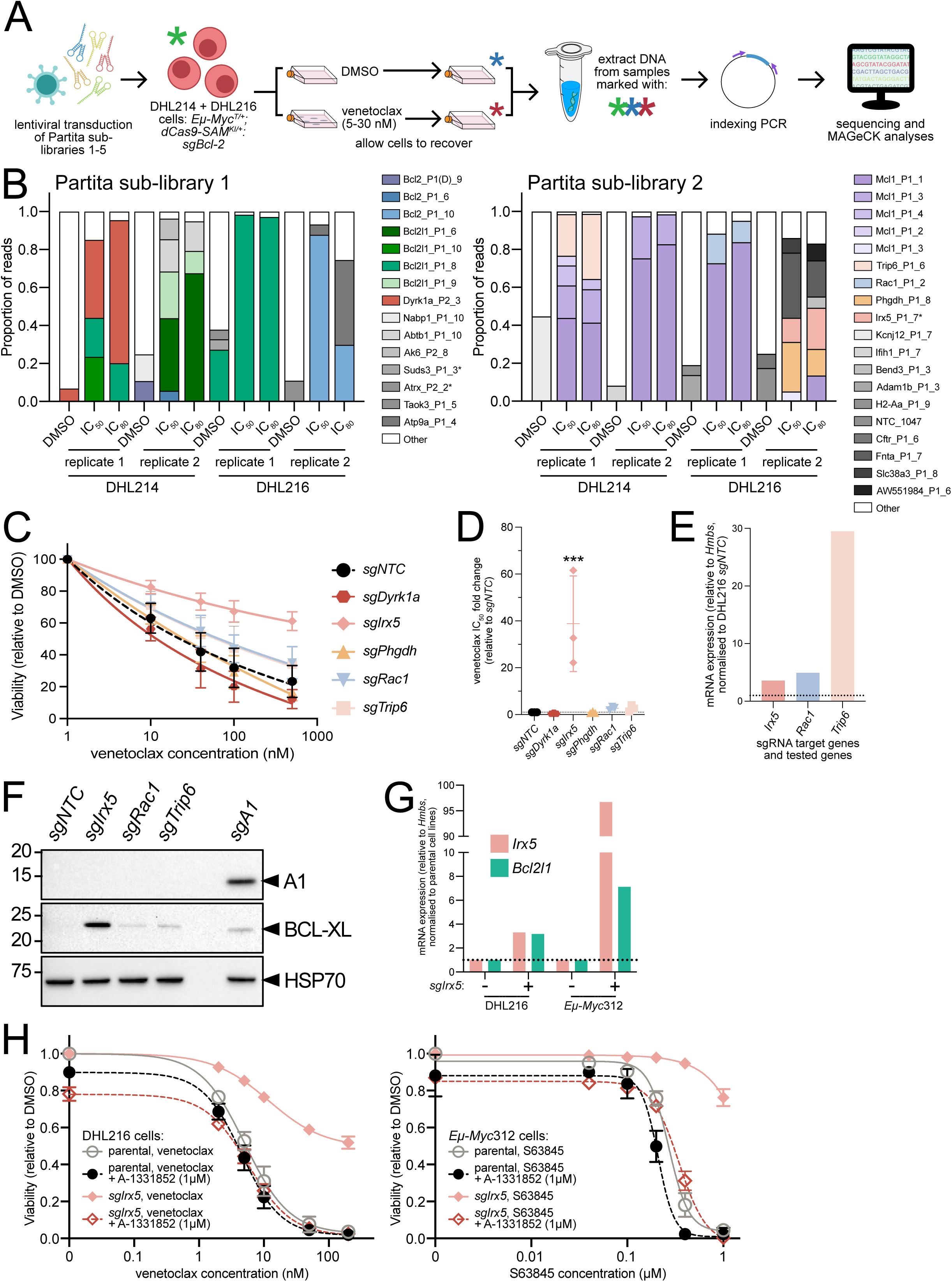
Whole genome screening to identify novel resistance factors for venetoclax treatment in lymphoma. (A) DHL cells (*Eµ-Myc^T/+^;dCas9a-SAM^KI/+^:sgBcl2*) were separately transduced with Partita sub-libraries 1-5 and then rapidly expanded. ∼10 days post-transduction, “input” cell pellets were collected, and DHL214/DHL216 cells were then treated with DMSO, or with venetoclax at concentrations sufficient to kill ∼90% (or more) of cells (n=2-3 independent treatments per library-transduced cells). After recovering, the cells were collected for DNA extraction, and then indexing PCR and NGS was performed. (B) Graphs showing enrichment of sgRNAs in DHL214/216 cells transduced with sgRNA sub-libraries 1 and 2 and treated with DMSO (control) or venetoclax. The x-axis indicates the different treatment conditions and replicates for each. Those sgRNAs indicated in the legend represent those with a count frequency of ≥5%. Certain known genes or potential genes of interest are shown in brighter colours (non-grey). Results for sub-libraries 3-5 are shown in Fig S3C. (C) Viability assay to validate the effect of enhanced target gene expression on cell survival over 24 h of venetoclax treatment (n=3 biological replicates per sample, each with 2 technical replicates per treatment). DHL216 cells were transduced with individual sgRNAs and treated with increasing concentrations of venetoclax. (D) Quantification of the IC_50_ values for venetoclax in DHL216 cells transduced with individual sgRNAs for each of the candidate genes (from C). Transduction of cells with *sgIrx5* led to a significant increase in IC_50_ (one-way ANOVA with Dunnett’s post-hoc test, p=0.0005). (E) qRT-PCR measuring gene expression of candidate genes in DHL214 cells transduced with individual sgRNAs targeting each gene (n=1 biological replicate per sample). Gene expression is measured relative to the housekeeping gene *Hmbs* and normalised to expression in the DHL216 cell line transduced with *sgNTC*. (F) Western blot showing expression levels of pro-survival proteins BFL-1/A1 and BCL-XL in DHL216 cells transduced with individual sgRNAs for each of the candidate genes. Probing for HSP70 was used as a loading control. DHL214 cells containing an sgRNA targeting BFL-1/A1 were used as a positive control. Additional western blots are included in Fig. S3D. Uncropped western blots are provided in Supplementary File 2. (G) qRT-PCR measuring the expression of *Bcl2l1* (encodes BCL-XL) and *Irx5* in DHL216 and *Eμ-Myc312* cells transduced with either *sgNTC* or *sgIrx5* (n=1 biological replicate per sample). *Bcl2l1* is transcriptionally induced by *Irx5* overexpression. Gene expression is measured relative to the housekeeping gene *Hmbs* and normalised to expression in each cell line transduced with *sgNTC*. (H) Viability assays to examine the ability of BCL-XL inhibitor A-1331852 to rescue resistance to the BCL-2 inhibitor venetoclax or the MCL-1 inhibitor S63845 in DHL216 or *Eµ-Myc312 sgIrx5* cells, respectively (n=3 biological replicates per sample, each with 2 technical replicates per treatment). Treatment with the BCL-XL inhibitor A-1331852 resensitised the *Irx5*-overepxressing cells to venetoclax or S63845. Where relevant, error bars represent mean ± SD. *** = p<0.001.

In parallel, we also conducted two additional screens using etoposide (a DNA damaging agent) or nutlin-3a (an MDM2-inhibitor), which both induce apoptosis through activation of TRP53 (*34, 35*) (Fig. S4A). We transduced just the DHL214 cell line with sub-libraries 1-5 (8-15% BFP- positive cells), again followed by expansion for ∼10 days. Cells were treated with drug concentrations that killed ≥90% of parental cells (Table S1) or with DMSO. The cells were then allowed to recover before DNA collection and sequencing (Fig. S4A).

In the venetoclax screen, coverage for both sgRNAs and genes was high in the input samples despite low initial fold library coverage (∼5-fold), with ∼90% of sgRNAs detected in cells prior to treatment (Fig. S3B). We next calculated the proportions of each sgRNA within each DMSO and venetoclax-treated sample (Figs. 3B, S3C; count data available at GSE296642). We observed sgRNAs targeting *Bcl2*, *Mcl1*, and *Bcl2l1* becoming enriched after venetoclax treatment, as would be expected from the well-characterised pro-survival roles of their encoded proteins (*8, 36*). We selected five novel hits to pursue for validation based on increasing proportions of sgRNAs targeting these genes in venetoclax-treated samples compared to DMSO-treated cells, as well as relevant cancer-links in published literature: *Irx5*, *Rac1*, *Trip6*, *Dyrk1a*, and *Phgdh* (Table S2). Individual sgRNAs for each gene were cloned and transduced into DHL216 cells, alongside a non-targeting control sgRNA (sgNTC) (Table S4). Viability assays over 24 h treatment with venetoclax at varying concentrations (though acute treatments and not fully representative of the longer term screening treatments) revealed that while expression of sgRNAs targeting *Rac1* and *Trip6* resulted in slight increases in venetoclax IC_50_ values, only targeting of *Irx5* shifted the venetoclax IC_50_ in DHL216 cells to a statistically significant extent (Fig. 3C,D). qRT-PCR confirmed that *Rac1*, *Trip6*, and *Irx5* were indeed upregulated by their corresponding sgRNAs (Fig. 3E).

Similarly high gene coverage was obtained for the nutlin-3a and etoposide screen input samples with >95% of genes being covered by at least one sgRNA (Fig. S4B). For these screens, we used MAGeCK (*37*) to identify sgRNAs enriched in the various treated samples compared to the DMSO control (Fig. S4C; count data available at GSE296642). As for the venetoclax screen, our approach was again validated by the identification of *sgBcl2* as the top hit from sub-library 1. The top ten most-enriched genes for each library and treatment were then identified via significance score (false discovery rate). The hits were further stratified based on: the number of independent sgRNAs identified per gene, gene/sgRNA enrichment across multiple replicates, gene/sgRNA enrichment across multiple treatments, and literature searches for logical candidates. According to these criteria nine genes were chosen for independent validation experiments: *Rnf128*, *Hira*, *Aoc2*, *Pik3c2a*, *Hic2*, *Ssxb10*, *Phgdh* (also a hit in the venetoclax screen), *Dnaja1*, and *Gsdma* (S Table 3). DHL214 transduced with individual sgRNAs targeting each hit (Table S4) had their gene induction confirmed by qRT-PCR, although variable induction levels were observed (Fig. S5A). We then subjected each cell line to 24 h viability assays with a range of concentrations of nutlin-3a or etoposide. The *sgDnaja1* validated in the former treatment, whereas *sgPik3c2a* validated in the latter, and *sgHira* and *sgPhgdh* validated in both treatments (Fig. S5B, Table S3). However, no significant shift in IC_50_ values were observed in any instance (Fig. S5C). To better reflect the chronic treatments from the screening protocol, 14 day cell competition assays were performed (for both nutlin-3a and etoposide) to tease out differences in sgRNA-driven survival that may only manifest over longer periods of treatment. These long-term assays revealed some similarities, as well as differences compared to the 24 h cell viability assay results (Fig. S5D, Table S3). *sgHira* and *sgDnaja1* transduced cells strongly outcompeted *sgNTC* transduced cells in both nutlin-3a and etoposide conditions (with *sgDnaja1* cells also exhibiting a growth advantage in DMSO (control) conditions). *sgAoc2*, *sgPhgdh*, and *sgPik3c2a* provided a competitive advantage in etoposide conditions only (the latter only very weakly). The *sgGsdma*, *sgHic2*, *sgRnf128*, and *sgSSxb10* did not validate under either condition tested. Interestingly, *Hira*, *Dnaja1*, and *Phgdh* have all been reported to encode modulators of TP53/TRP53 pathway activity. *PHGDH* repression by activated TP53 is necessary to promote apoptosis in certain contexts (*38*); DNAJA1 has been implicated in promoting metastasis by preventing mutant TP53 degradation (*39*); and dominant-negative HIRA prevents senescent behaviour driven by TP53 (*40*). While PIK3C2A has not been closely linked to this type of oncogenic potential, phosphatidylinositol 3-kinases (PI3Ks) are well known to have roles in the development and expansion of cancers (*41*). Those of the class 3 type (e.g. PIK3C2A) are the least well-understood members of this family. However, TP53 has been shown to transcriptionally regulate PIK3CA levels to effect apoptosis in some cancer cells (*42*).

As *Irx5* represented the strongest hit from all three screens overall, we chose to characterise its function in more detail. As the mechanism of action of venetoclax is induction of apoptosis, we performed western blotting for key intrinsic apoptosis pathway regulators (Figs. 3F, S3D). In cells upregulating *Irx5* (as well as those transduced with either *sgNTC*, *sgRac1*, or *sgTrip6*), we observed no obvious changes in the expression of pro-survival proteins BCL-2, MCL-1, or A1 (BFL-1 in humans), nor in the pro-apoptotic proteins PUMA and BIM, nor in TRP53 or its regulator P19ARF (Figs. 3F, S3D). However, we did observe a striking upregulation of BCL-XL in the DHL216 *sgIrx5* cells, indicating that this pro-survival factor may be the driving force behind this resistance to venetoclax (Fig. 3F). To determine whether induction of *Irx5* was transcriptionally upregulating BCL-XL, we performed qRT-PCR for *Bcl2l1* (which encodes BCL-XL) in DHL216 cells transduced with *sgIrx5* (Fig. 3G), as well as similarly transduced *Eμ-Myc*312 pre-B/B lymphoma cell line. We found that *Bcl2l1* was indeed transcriptionally induced in response to *Irx5* upregulation.

To confirm whether the increased BCL-XL in these cells was protecting them from apoptosis induced by venetoclax, we undertook viability assays using either parental or *sgIrx5*-transduced DHL216 cells in the presence of venetoclax alone or in combination with the BCL-XL-inhibiting BH3-mimetic drug A-1331852. Interestingly, we found that targeting of BCL-XL resensitised the *Irx5*-overexpressing cells to venetoclax, suggesting that venetoclax resistance in these cells was due to compensatory pro-survival expression of BCL-XL (Fig. 3H). To investigate whether overexpression of *Irx5* could also cause resistance to other BH3-mimetic drugs, we treated *Eμ- Myc*312 *sgIrx5* lymphoma cells with S63845 (a BH3-mimetic tool compound that specifically inhibits MCL-1, the pro-survival protein upon which *Eµ-Myc* cells depend (*43, 44*)). Similar to the results with venetoclax in DHL cells, the *Eμ-Myc*312 cells overexpressing *Irx5* were more resistant to S63845, but were resensitised to S63845 after targeting BCL-XL with A-1331852 (Fig. 3H). Overall, these data identify *Irx5* as a novel and potent resistance factor for BH3-mimetic drug therapy in blood cancers, demonstrating the power of these new CRISPR libraries to identify novel drug resistance factors.

These data demonstrate our new CRISPRa sgRNA libraries can be employed to explore biologically interesting questions and identify known and novel modulators of TRP53-mediated apoptosis in difficult-to-transduce mouse-derived cell lines.

### Screening in primary cells in vivo to uncover oncogenic factors in Myc-driven lymphomagenesis

Lastly, we aimed to demonstrate utility of the Partita library in primary cells during disease development *in* vivo. We used non-malignant primary haematopoietic stem and progenitor cells (HSPCs) sourced from the foetal livers of *Eµ-Myc* transgenic E14.5 embryos that, when transplanted into recipient mice, will spontaneously generate pre-B/B cell lymphoma (*30, 31*). By performing a pooled *in vivo* CRISPRa screen we anticipated identifying oncogenes that co- operate with MYC overexpression to accelerate B cell lymphoma development, similar in design to *in vivo* CRISPR knock-out screens previously performed by our lab (*45, 46*). In addition to using Partita sub-libraries one-five covering the whole mouse genome, we also utilised the transcriptional modifier-specific sub-library six, with the goal of identifying potential oncogenic transcription factors involved in lymphomagenesis.

To this end, *Eµ-Myc^T/+^;dCas9a-SAM^KI/+^* primary foetal liver cells (FLCs) were separately transduced with sub-libraries one-six (hereafter referred to as Partita1-6) (Fig. 4A). Cell transduction efficiency ranged between 17-40%, as determined by BFP fluorescence (Fig. S6A,B). Transduced FLCs were transplanted into lethally irradiated congenic *C57BL/6-Ly5.1* wild-type recipient mice and monitored for lymphoma development (Fig. 4A). All recipient mice, regardless of sgRNA-transduced donor FLCs, developed characteristic *Eµ-Myc* lymphoma pathology (*30, 31*), as evidenced by high white blood cell and low platelet counts in peripheral blood, as well as enlarged lymphoid tissues (e.g. spleen, thymus, lymph nodes) (Fig. S7A,B). Recipient mice transplanted with *sgNTC*-transduced *Eµ-Myc^T/+^;dCas9a-SAM^KI/+^* FLCs developed lymphoma with a median latency of 125.5 days post-transplantation, while positive control mice transplanted with *sgBcl2*-transduced *Eµ-Myc^T/+^;dCas9a-SAM^KI/+^* FLCs developed lymphoma significantly faster, with a median latency of 72 days post-transplantation (Fig. 4B). Some mice transplanted with Partita1-6 transduced *Eµ-Myc^T/+^;dCas9a-SAM^KI/+^* FLCs also displayed accelerated disease onset, with a median latency of 84.5 days post-transplantation (Figs. 4B, S7C), suggesting that sgRNAs within the Partita sub-libraries induced gene expression capable of accelerating MYC-driven lymphomagenesis. Interestingly, mice transplanted with FLCs transduced with the transcriptional modifers sub-library (Partita6) displayed the fastest lymphoma onset (median latency of 54.5 days post-transplantation). This suggested various transcriptional modifiers were likely important drivers of MYC-driven lymphomagenesis.

**Figure 4.**
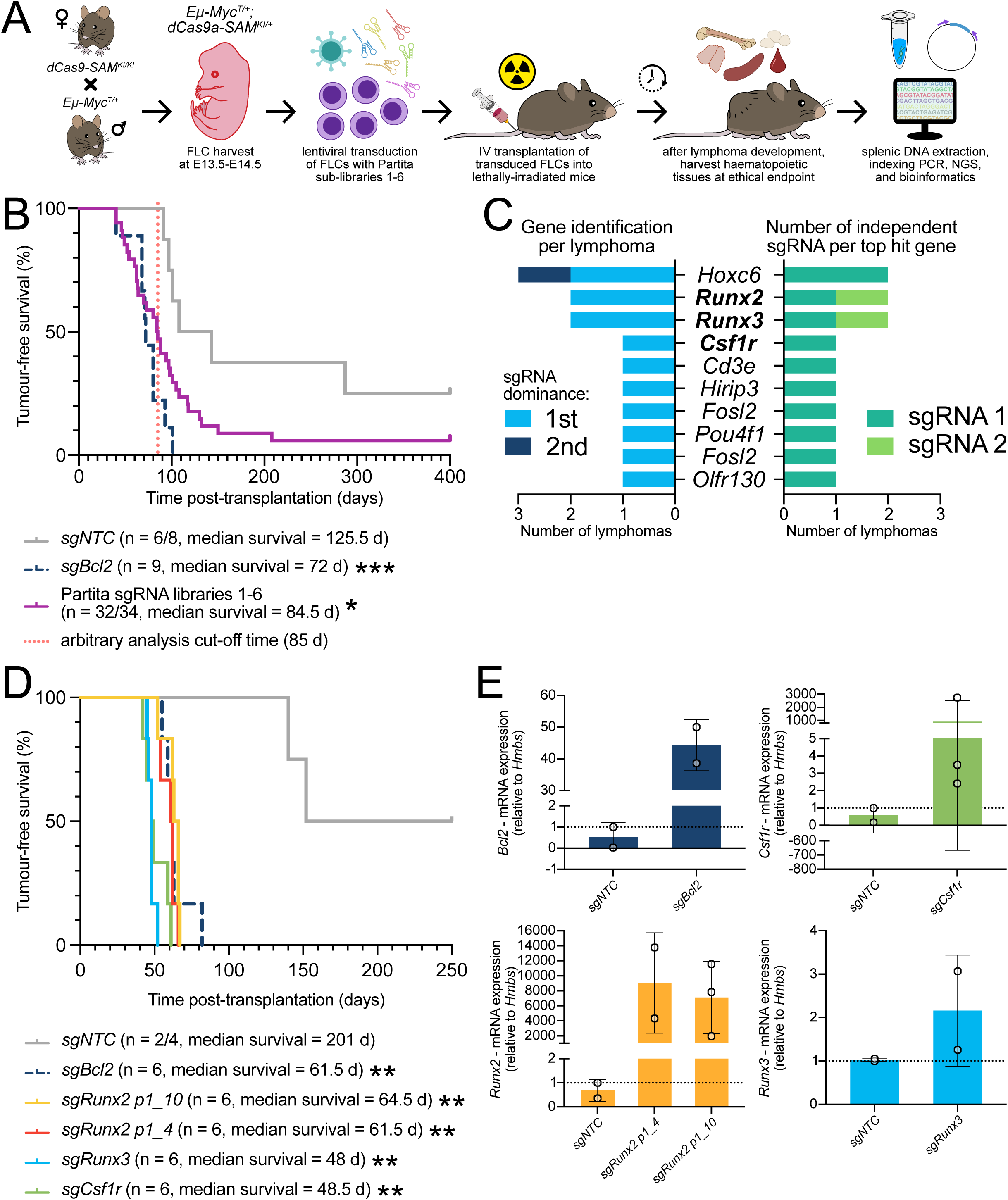
Employing novel sgRNA libraries in a pooled *in vivo* CRISPR activation screen to identify oncogenes that accelerate MYC-driven pre-B/B cell lymphomagenesis. (A) Schematic of *in vivo* pooled CRISPR activation screening approach employing the Partita genome-wide CRISPRa sub-libraries 1-5, as well as sub-library 6 (a boutique transcriptional modifier sub-library), using a haematopoietic reconstitution model system of wild-type mice transplanted with *Eµ-Myc^T/+^;dCas9a-SAM^KI/+^* foetal liver cells (FLCs; an abundant source of haematopoietic stem and progenitor cells (HSPCs)). This system enables the identification of genes that, when overexpressed, accelerate MYC-driven lymphoma development. (B) Tumour-free survival curve of lethally-irradiated recipient mice that had been transplanted with *Eµ-Myc^T/+^;dCas9a-SAM^KI/+^* FLCs that had been transduced with Partita1-6 (n=4-9 mice for each sub-library; see Fig. S7C for individual survival curves), or with a positive (*sgBcl2*, n=9) or negative (*sgNTC*, n=8) control guide. Recipient mice transplanted with Partita sub-library-transduced FLCs are combined into a single curve. A red dotted line indicates the cut-off time of 85 days post-transplantation used as an arbitrary indicator of accelerated lymphomagenesis. The data are a combination of two independent transplantation cohorts. The number of mice in each combined cohort are indicated, as are the median survival times. Statistical significance was measured by Mantel-Cox tests, comparing the *sgNTC* cohort to the *sgBcl2* (p=0.0006) and Partita1-6 (p=0.0114) cohorts. (C) Top 10 candidate genes identified in the *in vivo* screen. Candidate genes were ranked by: (i) their sgRNA dominance amongst detected reads (>20% of reads) in independent lymphomas from mice with accelerated lymphomagenesis, and (ii) the number of independent sgRNAs for a gene that are the most dominant within a lymphoma sample. Notably, *Runx2*, *Runx3*, and *Csf1r* (indicated in bold) belong to the same family of transcription factors. The left chart indicates the number of lymphoma samples in which each gene was the top or second top hit. The right chart indicates the number of lymphoma samples in which independent sgRNAs were identified as being enriched. (D) Tumour-free survival curve of lethally-irradiated recipient mice that were transplanted with *Eµ-Myc^T/+^;dCas9a-SAM^KI/+^* FLCs that had been transduced with individual sgRNAs targeting the top hits of *Runx2, Runx3*, and *Csf1r* (n=6 mice per sgRNA). Two sgRNAs were used for *Runx2* (*Runx2 p1_4* and *Runx2 p1_10*), and one for each of the other genes. Recipient mice that were transplanted with FLCs that had been transduced with positive (*sgBcl2*, n=6) or negative (*sgNTC*, n=4) control guides were also used. Statistical significance was measured by Mantel-Cox tests, comparing the *sgBcl2* (p=0.0038), *sgRunx2.4* (p=0.0040), *sgRunx2.10* (p=0.0040), *sgRunx3* (p=0.0038), and *sgCsf1r* (p=0.0038) cohorts to the *sgNTC* cohort. (E) qRT-PCR data comparing the relative expression of mRNA of target genes *Bcl2, Csf1r, Runx2*, and *Runx3* in lymphoma cell lines derived from the recipient mice transplanted with *Eµ-Myc^T/+^;dCas9a-SAM^KI/+^* FLCs. Expression was measured relative to the housekeeping gene *Hmbs* and then normalised to gene expression in the *sgNTC* negative control cohort. Data is presented as the mean, with error bars representing ± SD, and each dot representing an independent cell line (n=2-3 per genotype). * = p<0.05, ** = p<0.01, *** = p<0.001.

sgRNA enrichment in lymphomas was used to prioritize candidate oncogenes for validation. For this, gDNA was isolated from the tumour-burdened spleens of recipient mice that had developed lymphoma prior to 85 days post-transplantation (n=19/34; arbitrary cut-off for analysis), and NGS was performed to identify enriched sgRNAs (count data available at GSE296642). Many low read count sgRNAs were also detected, likely representing sgRNAs integrated into non-malignant splenocytes. In some lymphoma samples, more than one sgRNA accounted for >20% of the read counts, confounding which sgRNA was the main driver of accelerated lymphomagenesis. However, for most of the accelerated samples there were a small number of dominant sgRNAs (represented by >20% of sequencing read counts), which were the focus of our follow-up assays (Fig. S7D). We increased ranking of each candidate gene if its associated sgRNA was a strong hit in more than one accelerated sample, or if distinct sgRNAs targeting a gene were identified across multiple samples (Figs. 4C, S7D). It is important, however, to consider that spontaneous mutations in the TRP53 tumour suppressor pathway occur in ∼20% of *Eµ-Myc* lymphomas, and leads to strong acceleration of lymphomagenesis (*47–49*). If present, this would confound whether lymphomagenesis acceleration was due to the sgRNA(s) detected, or spontaneous mutations in TRP53. TRP53 pathway status was therefore assessed by western blotting to detect stabilised TRP53 protein and/or downstream P19 protein expression (increased upon TRP53 loss-of-function) (Fig. S7E). Hits identified in accelerated lymphoma samples with TRP53 pathway defects were excluded.

Overall, the strongest candidate genes were *runt-related transcription factor 2* (*Runx2*) and *Runx3*, with each having two different sgRNAs identified in two independent accelerated lymphoma samples (Fig. 4C). *Runx2* and *Runx3* belong to the same family of transcription factors known to be involved in normal haematopoiesis. Of note, overexpression of *RUNX2* and *RUNX3* has previously been implicated in the progression of both human lymphoid and myeloid malignancies (*50–52*). We also observed enrichment of *colony stimulating factor 1 receptor* (*Csf1r*), previously implicated in human blood cancers. The chromosomal rearrangement of this locus being observed in paediatric B cell acute lymphoblastic leukemia, and its expression correlating with a poor prognosis in follicular lymphoma (*53, 54*). Therefore, sgRNAs targeting *Runx2*, *Runx3*, and *Csf1r* were chosen for validation in independent FLC reconstitution assays to evaluate their potential for accelerating MYC-driven lymphomagenesis. Individual sgRNAs targeting these genes (1 sgRNA for each of *Runx3* and *Csf1r*, 2 sgRNAs for *Runx2*; Table S4), taken from the Partita6 library, were transduced into *Eµ-Myc^T/+^;dCas9a-SAM^KI/+^* FLCs that were then transplanted into lethally-irradiated wild-type congenic recipient mice (alongside *sgNTC* or *sgBcl2* negative and positive controls, respectively). The FLC-transplanted mice were then monitored for lymphoma development. Each of these sgRNAs independently validated as accelerants of MYC-driven lymphomagenesis (Fig. 4D), with all recipient mice displaying characteristic lymphoma pathology at endpoint (Fig. S8A-D). In cell lines derived from tumours from these mice, transactivation of each target gene was confirmed by qRT-PCR. Relative to the *sgNTC* samples, a clear increase in target gene expression was observed for each sgRNA of interest (Fig. 4E). These findings demonstrate the utility of the Partita sgRNA sub-libraries as tools to discover and investigate genes involved in lymphoma development and for *in vivo* screening applications.

## Discussion

CRISPR activation has for several years now enabled biological studies from a new point-of-view. In this study, we describe the creation of a new whole-genome CRISPRa sgRNA library – Partita. Partita is split into five sub-libraries targeting different classes of genes, as well as a sixth sub-library only targeting transcriptional modifiers. We further validate Partita through a variety of screening approaches: enrichment/depletion screens in iBMDMs, *ex vivo* screens to identify anti-cancer drug resistance factors in a DHL model, and *in vivo* screening to discover co- operative mutations in lymphoma development.

There are a number of murine CRISPRa sgRNA libraries already available for a variety of CRISPRa systems (*2*), including (to our knowledge) two compatible with the *dCas9-SAM* model: mouse SAM v1 (*3, 12*) and Caprano (*13*). These whole-genome libraries have proven useful to the scientific community, having been applied by ourselves (*8*) and many other researchers to a wide variety of biological questions. We have designed Partita to build upon the advantages these libraries have provided, while also improving upon the high standard that already exists. For example, mouse SAM v1 uses three sgRNAs per gene, and Caprano use six sgRNAs per gene. Partita employs 10 scaffold-optimized (*16*) sgRNAs for the majority of the genome; higher density targeting with additional sgRNAs is designed to increase the probability that at least one will be capable of robust gene activation for each locus. Going beyond this, we have included sgRNAs intended to activate ≥1 TSS per gene, to account for tissue- or isoform-specific effects of transcripts expressed from distinct promoters. We have chosen to divide Partita into five unique sub-libraries. While mouse SAM v1 has no sub-libraries, a similar approach was taken with Caprano, where the six sgRNAs per gene are divided into two roughly equally sized sub-libraries (therefore three sgRNAs per gene per sub-library). We believe the five sub-libraries of Partita may prove advantageous for certain experimental approaches, helping to achieve higher library coverage, particularly important given the higher number of sgRNAs per gene. The sub-libraries may also make it possible for users to more easily target certain groups of genes, given we have divided them based on distinct classifications. Additionally, both mouse SAM v1 and Caprano use puromycin as a selection marker. In the Partita vector backbone we have included both puromycin and BFP, which we hope will aid certain users or in methodologies.

We have demonstrated the efficacy and utility of Partita in each of our screening contexts. In each instance, technical success of the library and screen execution was shown by our ability to recover a variety of well-characterized targets for each experimental context. For example, the iBMDM enrichment/depletion screen findings supported the known function of several pro-survival genes, most prominently the BCL-2 family member *Bcl2l1* (encoding BCL-XL), while also verifying the growth-inhibitory effects of activating certain genes that promote cell death or senescence, most prominently *Ripk1* and *Cdkn1a* (encoding P21), respectively.

In our screens to identify drug resistance modifiers in DHL cells, we anticipated and indeed observed the enrichment of sgRNAs targeting the pro-survival genes *Bcl2*, *Mcl1*, and, again, *Bcl2l1*. These are all well-characterised as promoting resistance to apoptosis-inducing anti- cancer agents in various blood cancers (*8, 36*). Thus, our results represent effective validation of our screen design and the Partita library. *Irx5* was a notable hit to come out of these screens; its upregulation conferred potent resistance to venetoclax treatment. *Irx5* encodes the Iriquois homeobox 5 protein and, unsurprisingly for a homeobox transcription factor, is mostly implicated in a variety of developmental roles, most prominently in the eyes and heart (*55–57*). However, *Irx5* upregulation has also been linked to the upregulation of NFκB signalling in certain contexts (*58*). This is intriguing, as NFκB signalling has also been implicated in the direct upregulation of *Bcl2l1*/BCL-XL (*59*). As we demonstrated *sgIrx5* lymphoma cells have increased BCL-XL expression, and that they then show sensitivity towards treatment with the BCL-XL inhibitor A-1331852, it is plausible that this Irx5-NFκB-*Bcl2l1* axis may be what has been triggered in these assays. We are pursuing these questions as part of our future work. Irx5 has been studied in certain cancer contexts. *IRX5* knock-out hepatocellular carcinoma cell lines have been observed to have enhanced activation of the pro-apoptotic TP53 signalling pathway, coupled with reduced expression of BCL-2, though, unfortunately, BCL-XL expression was not examined in that study (*60*). In haematopoietic malignancies, overexpression of *IRX5* (as well as family members *IRX1* and *IRX3*) has been observed in some acute myeloid leukaemia cell lines, though the precise functional consequences of IRX5 in this context has not been elucidated (*61*).

We also identified several strong candidate genes in our *in vivo* screen to identify accelerants of *Myc*-driven lymphomagenesis. While we did not validate *Hoxc6* in the *Eµ-Myc* context individually, it is known that *Hoxc6* is overexpressed in various human lymphoid leukaemias (*62*), and that its ectopic expression in isolation is capable of driving myeloid malignancies (*63*). The genes we did successfully validate – *Runx2*, *Runx3*, and *Csf1r* – have also all been implicated in haematopoietic malignancies. Both *Runx2* and *Runx3* have been shown to be involved in several cancer contexts in a variety of different roles (*50*). *Runx2* has long been known to be involved in lymphomagenesis. Certain retroviral insertions were shown to be capable of driving *Runx2* expression (then called *Cbfa1*), and it was identified that this ectopic *Runx2* expression co-operated with *Myc* over-expression (*64, 65*), *Trp53* loss, or *Pim1* activation (*52*) to drive T cell lymphoma development. *Runx3* represents a similar situation, having also been shown to co- operate with *Myc* over-expression in T cell lymphomagenesis after Moloney murine leukemia virus insertion leads to *Runx3* upregulation (*66*).

*Csf1r* is best known for its well-established role as a key regulator of myeloid cell development, differentiation, and survival, particularly of monocytes and macrophages (*67*). Mice deficient for *Csf1r* lack the majority of their macrophage populations (*68*), while stimulation of Csf1r via its ligands Csf1 or IL-34 can promote macrophage proliferation (*69, 70*). To our knowledge, *Csf1r* has not previously been demonstrated to co-operate with *Myc* over-expression to promote tumourigenesis, but such a role is consistent with previous studies, where *CSF1R* has been implicated in numerous haematopoietic malignancies (*71*). For example, oncogenic fusion proteins containing CSF1R have been observed in cases of B cell precursor acute lymphoblastic leukaemia (B-ALL) (*72–74*) and acute megakaryoblastic leukaemia (*75*). Moreover, constitutively activating mutations in *CSF1R* have been associated with chronic myelomonocytic leukemia and AML (*76*). Targeted inhibition of *CSF1R* has been proposed as a potential therapeutic in AML, where its suppression prevents disease-supporting cells from releasing growth factors and cytokines (*77*), as well as in follicular lymphoma, where CSF1R inhibition promotes macrophage reprogramming and microenvironmental changes (*53*).

Gain-of-function screening technologies, like CRISPRa, enable unique discovery potential compared to loss-of-function methods, such as RNAi or CRISPR KO. For example, as we and others have demonstrated through passive growth screens, targeted gene induction can recapitulate phenotypes associated with locus amplification/aneuploidy, resulting in so-called “STOP” and “GO” effects (*78*). We have utilised Partita largely in the context of myeloid cell types and blood cell cancers, but, as with other pooled CRISPR screening platforms, it is designed to be applicable for a wide variety of experimental contexts and assays. For example, we have been particularly interested in exploring questions of fundamental biology using Partita sub-library six, which comprises transcriptional modifiers, to identify gene regulatory networks that contribute to haematopoietic lineage commitment during differentiation. It is likely that there are other gene class-specific libraries that could be designed based on the Partita sgRNA sequences. As CRISPRa matures, there is also significant interest in its potential therapeutic applications, particularly in the immunotherapy space. Screens looking to unravel gene regulatory networks in immune cells have been conducted previously (*79–81*), and such studies could potentially benefit from the new advantages that Partita brings to screening. Partita may be similarly useful in the advancement of newer technologies into the CRISPRa space, such as spatial transcriptomics which, to our knowledge, has so far only been combined with CRISPR knock-out methodologies (*82, 83*). Lastly, we and another group have recently published the first *Cas12a*-capable mouse models (*84, 85*). Cas12a is capable of being multiplexed with the dCas9-SAM system of CRISPRa, opening new avenues for the investigation of complex gene networks, or their modelling *in vivo*. We anticipate that Partita will find an increasingly expanded array of uses thanks to its power as a stand-alone or combinatorial screening platform.

## Supporting information

Partita Library Files

---

**Supplementary Figure 1.**
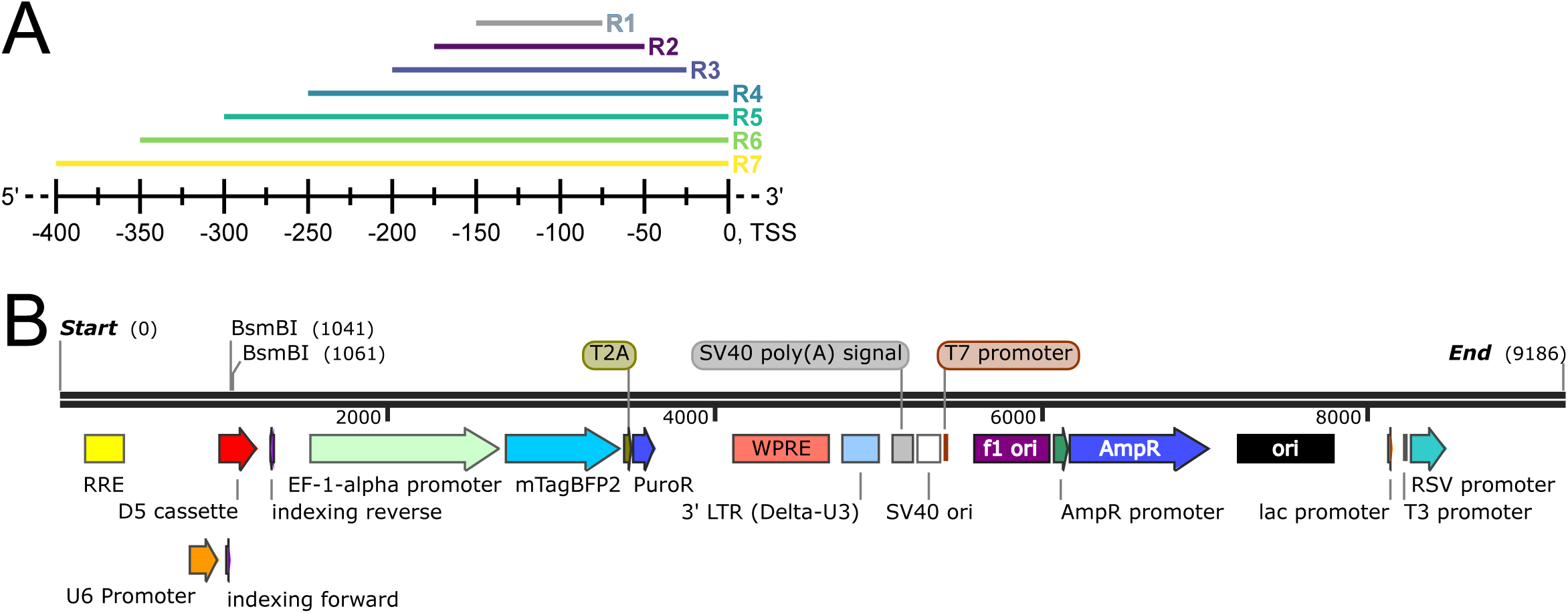
Details about the creation of the Partita library. (A) Diagram showcasing the prioritisation of sgRNA selection within the optimal activation window. Priority was given first to sgRNAs within the first region (R1) of [-150 bp, -75 bp], and then in regions of decreasing priority (and increasing size): R2 = [-175 bp, -50 bp], R3 = [-200 bp, -25 bp], R4 = [-250 bp, 0 bp], R5 = [-300 bp, 0 bp], R6 = [-350 bp, 0 bp], and R7 = [-400 bp, 0 bp]. (B) Plasmid map of the WEHI-12 vector used as the backbone for the Partita library. sgRNAs were cloned into the BsmBI cut site, within the D5 cassette, and under the control of the U6 promoter. A BFP reporter gene and a puromycin resistance gene under control of the EF-1α promoter are also included.

**Supplementary Figure 2.**
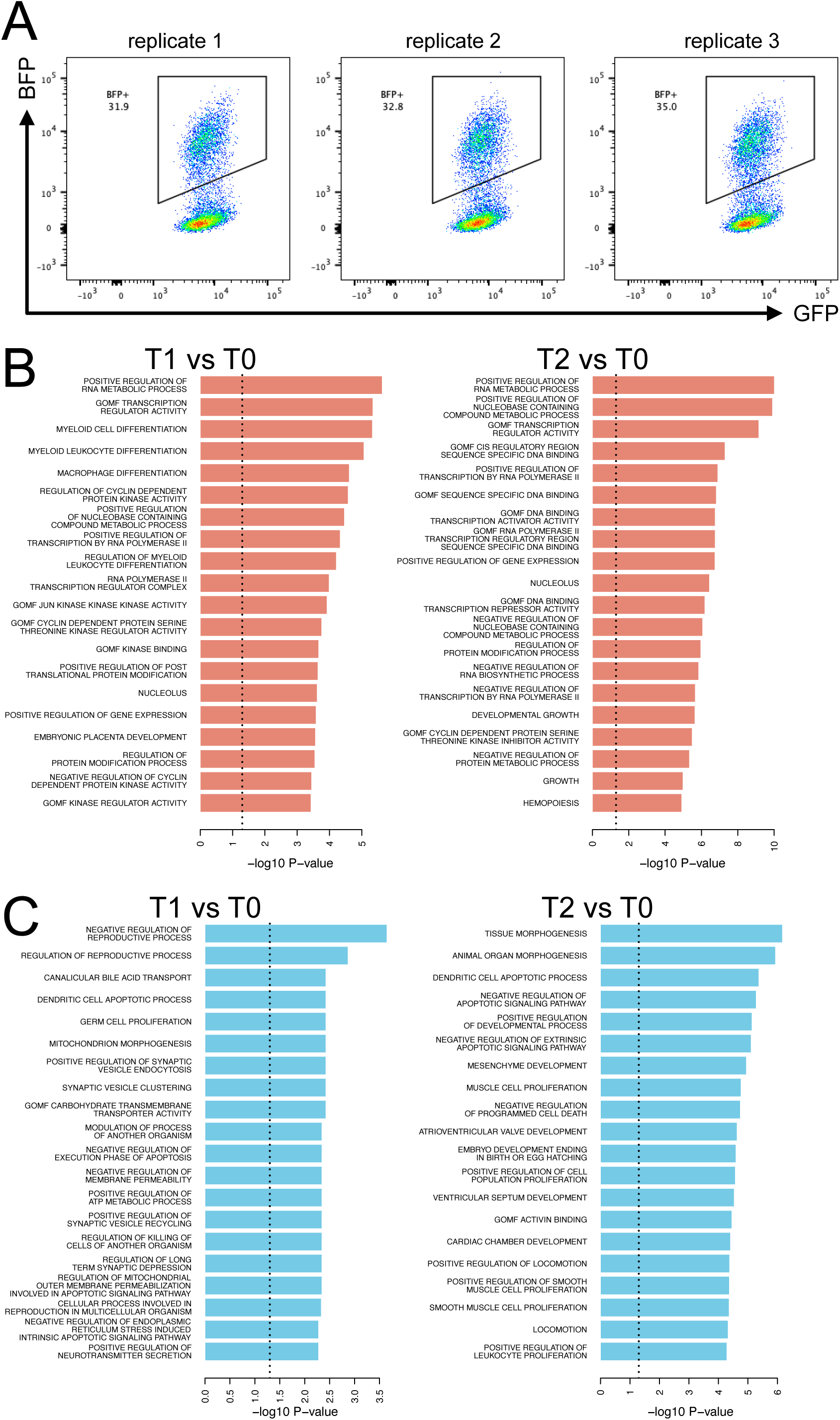
Additional data relating to the *in vitro* iBMDM enrichment/depletion screen. (A) Flow cytometry plots demonstrating the proportions of BFP-positive cells after lentiviral transduction of the iBMDMs with Partita sub-library 1. A 30-35% transduction efficiency was calculated as necessary to obtain one sgRNA per cell. (B) GSEA graphs showing the top twenty depleted gene classes at time points T1 and T2, relative to T0. Note the depletion of gene classes likely to be deleterious if activated, such as negative transcriptional regulators. Data for T3 vs T0 are shown in Fig. 2C. (C) GSEA graphs showing the top twenty enriched gene classes at time points T1 and T2, relative to T0. Note the enrichment of gene classes likely to be advantageous if activated, such as negative regulators of apoptosis. Data for T3 vs T0 are shown in Fig. 2D.

**Supplementary Figure 3.**
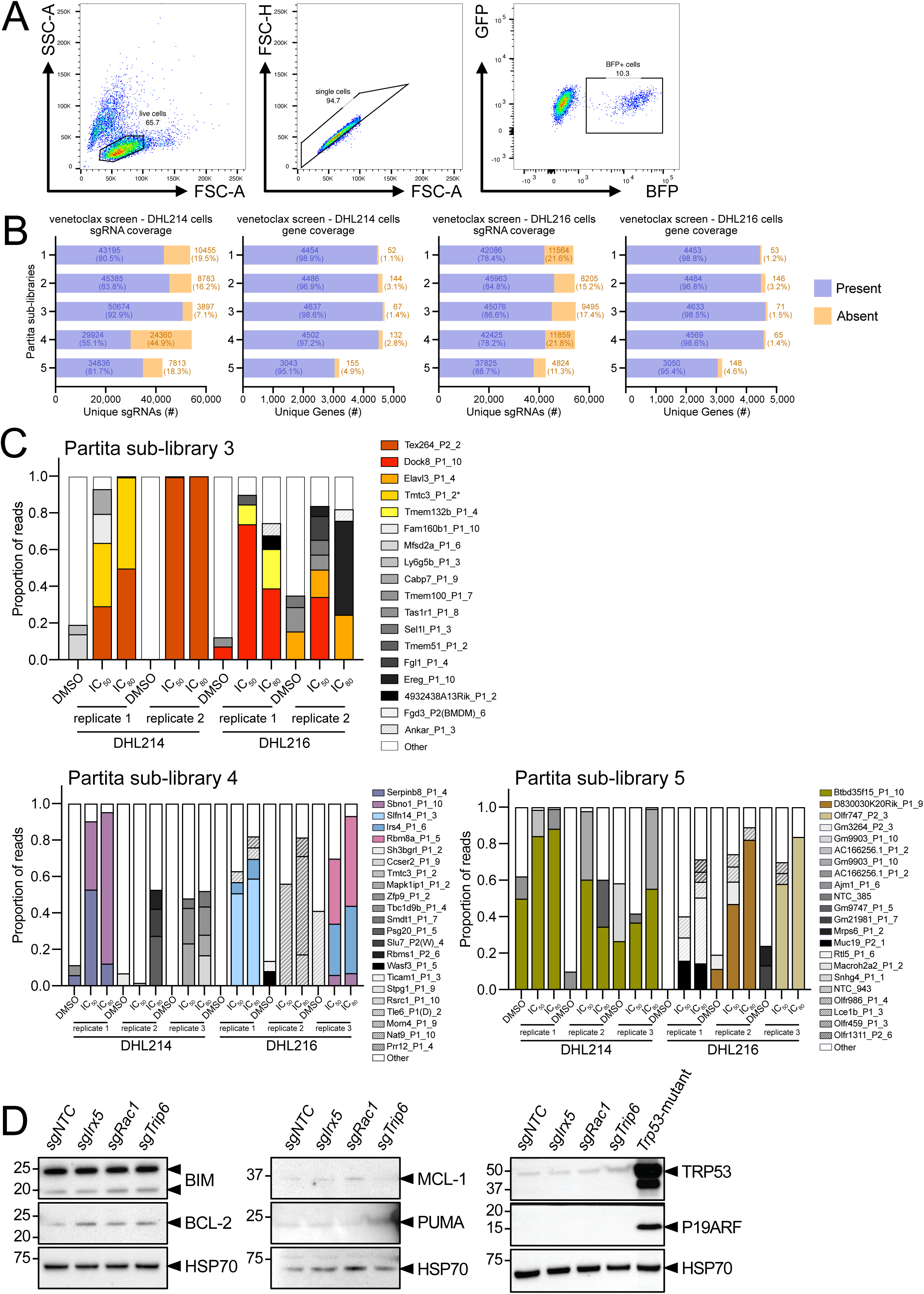
Additional data relating to *in vitro* drug resistance screening with venetoclax in lymphoma cells. (A) Representative flow cytometry gating of transduced DHL cells used in the venetoclax, nutlin-3a, and etoposide screens. Cells were gated first on live status and single cell status, based on forward and side scatter. Cells were then gated based on GFP-positivity (dCas9-SAM presence) and BFP-positivity (Partita sub-library presence). The example plots are for DHL216 cells transduced with sub-library 1. (B) Graphs indicating gene and sgRNA coverage for each sub-library in the DHL214 and DHL216 cells used in the screen with venetoclax. (C) Graphs showing enrichment of sgRNAs in DHL214/216 cells transduced with Partita sub-libraries 3-5 and treated with DMSO or venetoclax. The x-axis indicates the different treatment conditions and replicates for each. Those sgRNAs indicated in the legend represent those with a count frequency of ≥5%. Certain known genes or potential genes of interest are shown in brighter colours (non-grey). Results for sub-libraries 1 and 2 are shown in Fig. 3B. (D) Additional western blot indicating protein expression levels of various proteins in DHL216 cells transduced with individual sgRNAs for each of the candidate hit genes. HSP70 expression was used as a loading control. An additional *Trp53*-mutant sample was used as a positive control in the right-hand panel (this sample was a spontaneously *Trp53*-mutant *Eµ-Myc* lymphoma cell line).

**Supplementary Figure 4.**
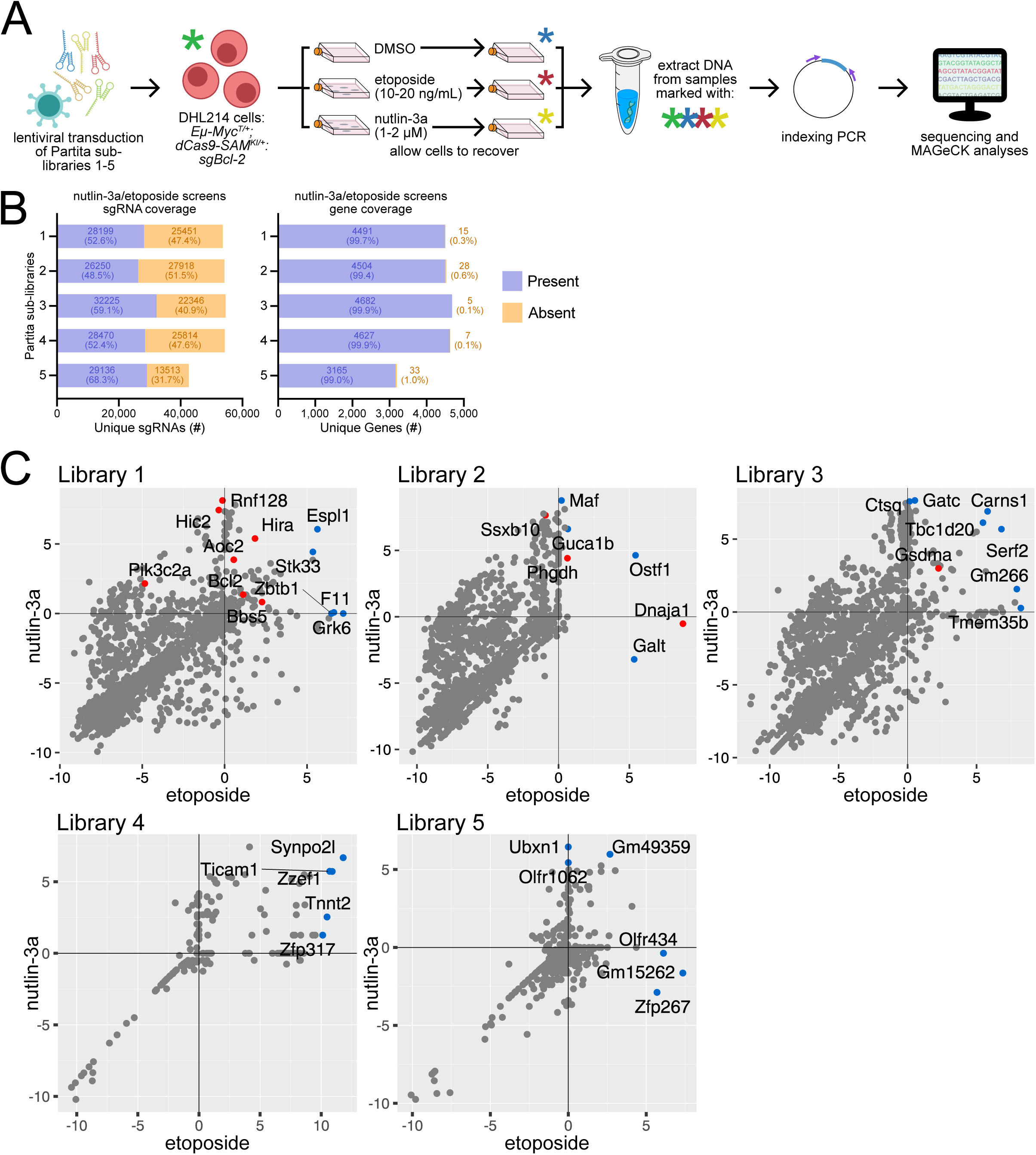
Whole-genome CRISPRa screening to identify regulators of TRP53-mediated apoptosis in lymphoma. (A) DHL cells (*Eµ-Myc^T/+^;dCas9a-SAM^KI/+^:sgBcl2*) were separately transduced with Partita sub-libraries 1-5 and then rapidly expanded. ∼10 days post-transduction, “input” cell pellets were collected, and DHL214 cells were then treated with DMSO (control), nutlin-3a, or etoposide, the latter two at concentrations sufficient to kill ∼90% (or more) of cells (n=3 independent treatments per library-transduced cells). After recovering, surviving cells were collected for DNA extraction, and then indexing PCR and NGS was performed. (B) Graphs indicating gene and sgRNA coverage in pre-treatment input samples for each sub-library in the DHL214 cells used in the screens with nutlin-3a and etoposide. (C) Plots showing log2 fold changes of genes in cells treated with nutlin-3a or etoposide for Partita sub-libraries 1-5, with each treatment measured relative to the DMSO control. Genes chosen for further validation are marked in red. Other genes with one or more sgRNAs significantly enriched, as determined by MAGeCK analyses, are marked in blue. All other genes are indicated in grey.

**Supplementary Figure 5.**
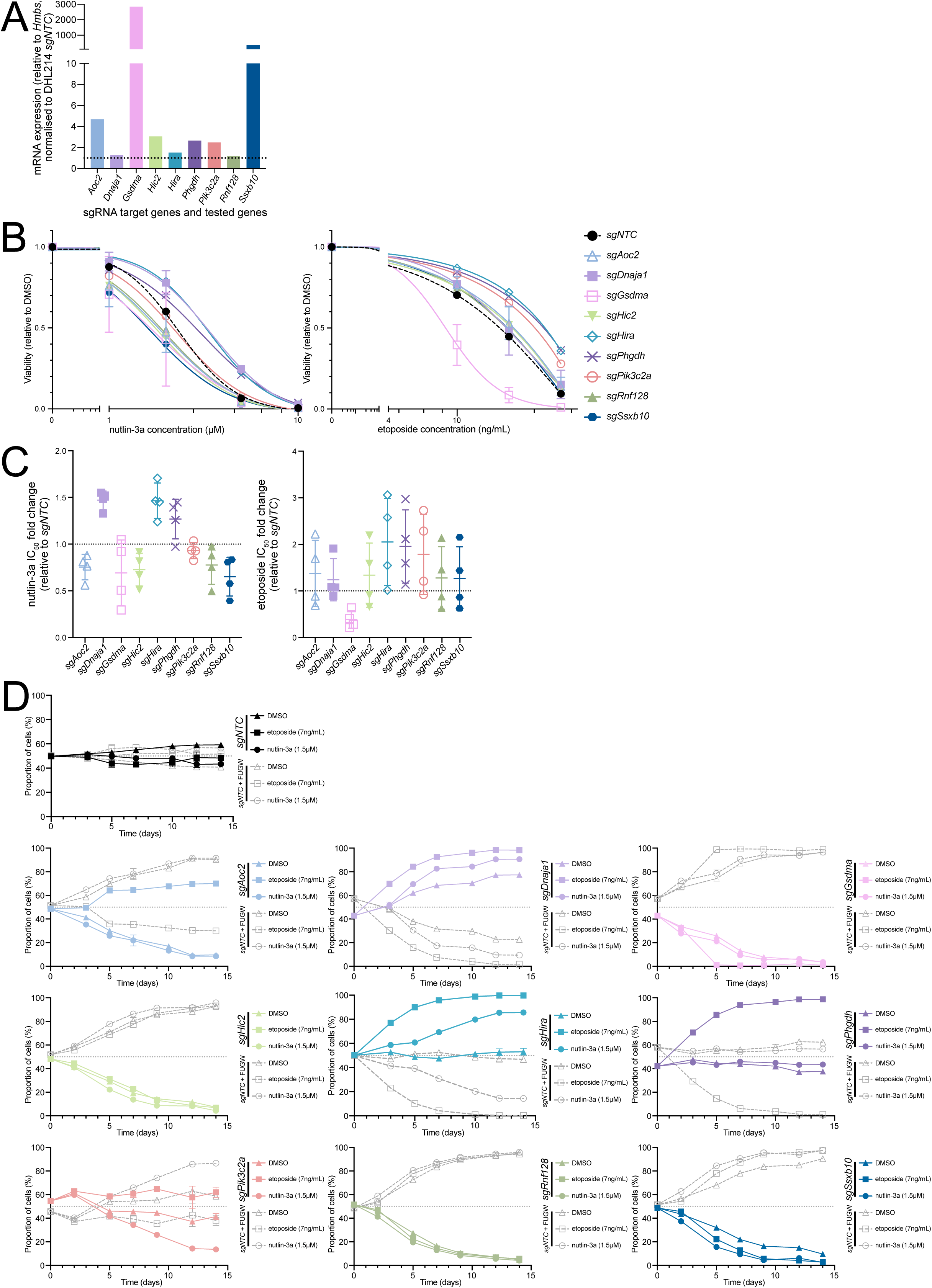
Additional data relating to *in vitro* drug resistance screening with nutlin-3a/etoposide in lymphoma cells. (A) qRT-PCR measuring gene expression of candidate genes in DHL214 cells transduced with individual sgRNAs targeting each gene (n=1 biological replicate per sample). Gene expression is measured relative to the housekeeping gene *Hmbs*, and normalised to expression in the DHL214 cell line transduced with *sgNTC*. (B) Viability assays to validate the effect of enhanced gene expression on cell survival over 24 h, in DHL214 cells transduced with individual sgRNAs (n=4 biological replicates per sample, each with 2 technical replicates per treatment). Cells were treated with increasing concentrations of either nutlin-3a or etoposide. (C) Quantification of the IC_50_ values for nutlin-3a and etoposide in DHL214 cells transduced with individual sgRNAs for each of the candidate genes (n=4 biological replicates per sample, each with 2 technical replicates per treatment). No changes were statistically significant (one-way ANOVAs with Dunnett’s post-hoc test), but some clear changes in IC_50_ values were still observed for some genes. (D) Cell competition assays *in vitro* used to validate the effect of enhanced gene expression on cell survival over 14 d, in DHL214 cells transduced with individual sgRNAs. Representative graphs are shown (n=2 independent assays per cell line, each with 2 technical replicates). Cells were treated with either DMSO (control), nutlin-3a (1.5 μM), or etoposide (7 ng/mL). The top graph shows competitiveness of *sgNTC*-transduced cells, and the remaining graphs show the competitiveness of cells transduced with individual sgRNAs targeting the various candidate genes. In all instances, competitiveness was measured in comparison to cells transduced with *sgNTC* (+ empty vector FUGW, which expresses strong GFP). Where relevant, error bars represent mean ± SD.

**Supplementary Figure 6.**
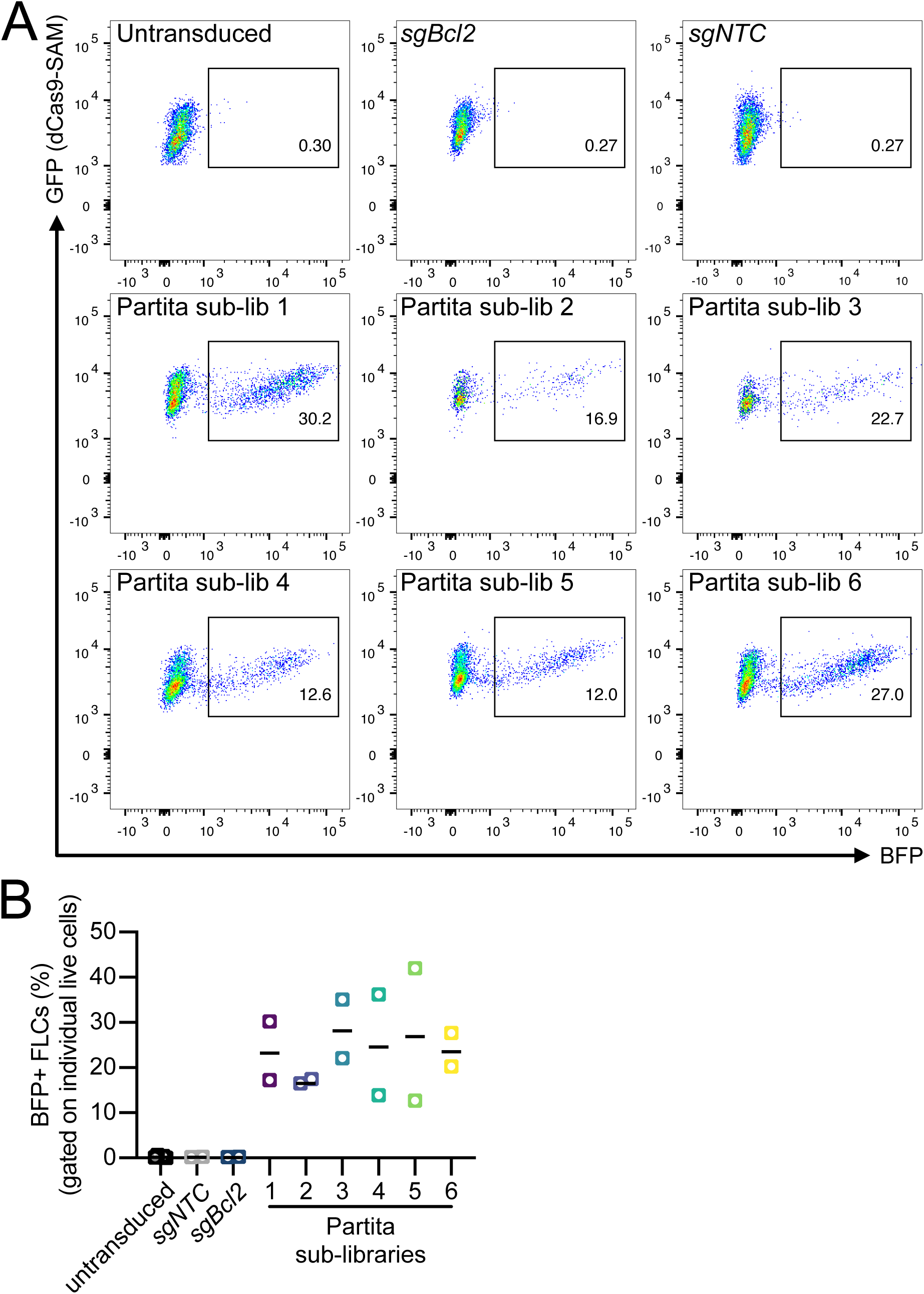
Additional data relating to the pooled *in vivo* CRISPRa screen. (A) Representative flow cytometry gating for *Eµ-Myc^T/+^;dCas9a-SAM^KI/+^* FLCs transduced with either the control vectors or Partita sub-libraries 1-6. The plots demonstrate the level of BFP detected in the FLCs, which can be used as a readout of transduction efficiency for the Partita library. (B) Graph quantifying the transduction efficiency (based on BFP status) for each of the different FLC lines used in the screen. Note that the non-transduced FLCs and those transduced with the control vectors had no BFP expression, as expected.

**Supplementary Figure 7.**
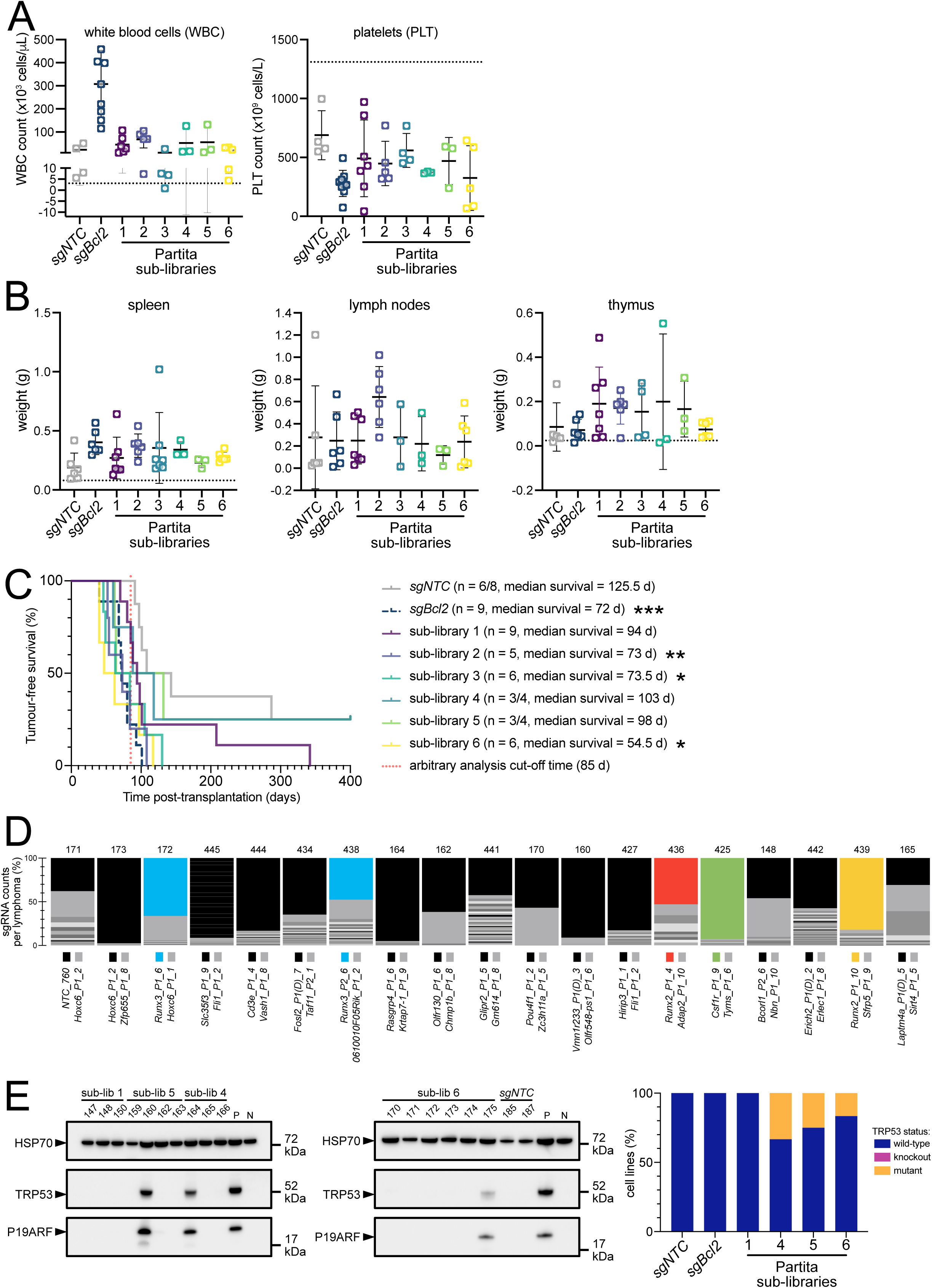
Additional data relating to the pooled *in vivo* CRISPRa screen. (A) Peripheral blood cells analyses from lethally-irradiated recipient mice that were transplanted with *Eµ-Myc^T/+^;dCas9a-SAM^KI/+^* FLCs that had been transduced with Partita1-6 (n=3-7 mice sgRNA cohort). From left to right, white blood cell (WBC) and platelet counts are shown, taken at ethical endpoint. Each data point indicates an individual mouse with mean ± SD. (B) Lymphoid organ analyses from lethally-irradiated recipient mice that were transplanted with *Eµ-Myc^T/+^;dCas9a-SAM^KI/+^* FLCs that had been transduced with Partita1-6 (n=3-7 mice sgRNA cohort). From left to right, the weights of the spleen, lymph nodes (inguinal, brachial and axial pairs), and thymus are shown, taken at ethical endpoint. Each data point indicates an individual mouse with mean ± SD. (C) Tumour-free survival curve of lethally-irradiated recipient mice that were transplanted with *Eµ-Myc^T/+^;dCas9a-SAM^KI/+^* FLCs that had been transduced with Partita sub-libraries, or with a positive (*sgBcl2*) or negative (*sgNTC*) control guide. Data are the same as those shown in Fig. 4B, but survival curves have been separated by sub-library cohort. The number of mice in each cohort are indicated, as are the median survival times. Statistical significance was measured by Mantel-Cox tests, comparing the *sgBcl2* (p=0.0006), Partita1 (p=0.0792), Partita2 (p=0.0039), Partita3 (p=0.0189), Partita4 (p=0.3431), Partita5 (p=0.3431), and Partita6 (p=0.0107) cohorts to the *sgNTC* cohort. (D) Visualisation of NGS data from the spleens of *Eµ-Myc^T/+^;dCas9a-SAM^KI/+^* FLC recipient mice with accelerated lymphoma (n=19). Tissues from each mouse were analysed by NGS to identify dominant sgRNAs and their targeted candidate oncogene that accelerates lymphomagenesis. Each bar graph represents one lymphoma sample, and each colour represents the percentage of counts for a sgRNA revealing its dominance. The two sgRNAs accounting for the greatest percentage of reads in each sample are indicated below each lymphoma. Genes with an sgRNA detected in more than one lymphoma are highlighted in colour. Lymphoma samples are ordered by lymphoma latency, fastest from left to right. Mouse numbers are indicated above each lymphoma sample. (E) Assessment of TRP53 protein wild-type or mutant status by western blotting for TRP53 and P19ARF. Blotting for HSP70 serves as loading control. The lane marked ‘P’ is a positive control mutant TRP53 *Eµ-Myc* lymphoma cell line with stabilised TRP53 and P19ARF protein expression, and the lane marked ‘N’ is a negative control wild-type TRP53 *Eµ-Myc* lymphoma cell line where wild-type TRP53 protein is maintained at low levels within a cell, suppressing P19 protein expression. A summary graph (right) indicates the proportion of different TRP53 protein statuses observed. * = p<0.05, ** = p<0.01, *** = p<0.001. Black dotted lines in A and B are representative phenotypic values from healthy male *C57BL/6* mice of either 10 (thymus) or 16 (all other cells/tissues) weeks old (*86–88*).

**Supplementary Figure 8.**
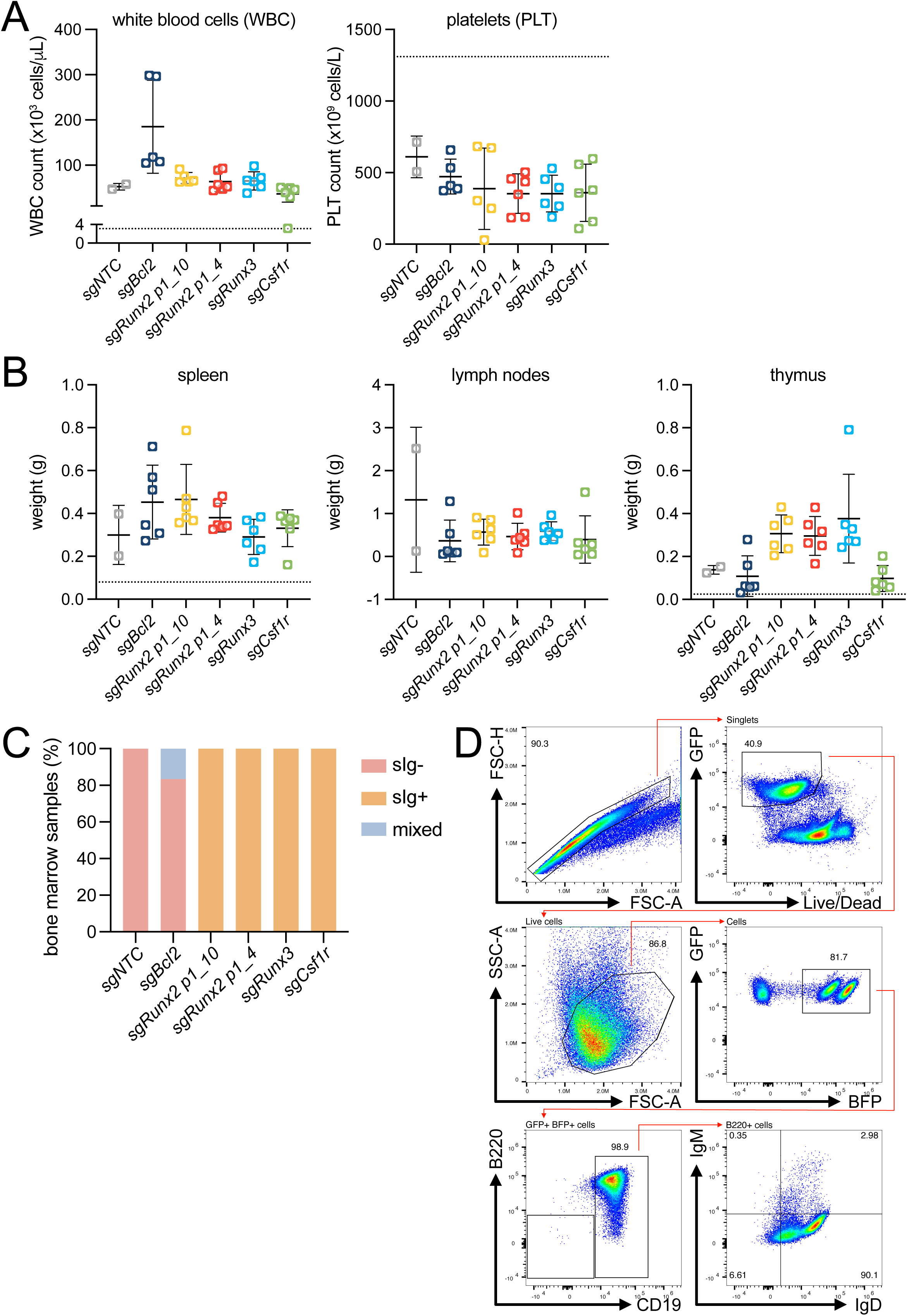
Additional data relating to validation of candidate oncogenes. (A) Peripheral blood cells analyses from lethally-irradiated recipient mice that were transplanted with *Eµ-Myc^T/+^;dCas9a-SAM^KI/+^* FLCs that had been transduced with individual sgRNAs (n=2-6 mice sgRNA cohort). From left to right, white blood cell (WBC) and platelet counts are shown, taken at ethical endpoint. Each data point indicates an individual mouse with mean ± SD. (B) Lymphoid organ analyses from lethally-irradiated recipient mice that were transplanted with *Eµ-Myc^T/+^;dCas9a-SAM^KI/+^* FLCs that had been transduced with individual sgRNAs (n=2-6 mice sgRNA cohort). From left to right, the weights of the spleen, lymph nodes (inguinal, brachial and axial pairs), and thymus are shown, taken at ethical endpoint. Each data point indicates an individual mouse with mean ± SD. (C) B cell lymphoma immunophenotyping data, as determined by flow cytometric analyses of bone marrow cells, taken at ethical endpoint from mice transplanted with *Eµ-Myc^T/+^;dCas9a-SAM^KI/+^* FLCs transduced with individual sgRNAs (n=2-6 bone marrow tissue samples per individual sgRNA cohort). Immunophenotypes were determined by staining with the B cell markers B220, CD19, surface IgM (sIgM) and IgD (sIgD). All lymphoma cells stained were B220 and CD19 double-positive, and either sIg- (both sIgM- and sIgD-; pro-B/pre-B), sIg+ (both sIgM+ and sIgD+; B), or sIg mixed (populations with both sIg- and sIg+ cells). (D) Representative flow cytometry gating for the immunophenotyping described in C. Black dotted lines in A and B are representative phenotypic values from healthy male *C57BL/6* mice of either 10 (thymus) or 16 (all other cells/tissues) weeks old (*86–88*).

## Materials and Methods

### Mouse husbandry and maintenance

Care and husbandry of experimental mice was performed according to the guidelines established by The Walter and Eliza Hall Institute Animal Ethics Committee, in accordance with the Australian Code of Practice for the care and use of animals for scientific purposes.

Mice were housed in a stabilised environment with 12-hour light and 12-hour dark cycles, with water and chow provided *ad libitum*. Transported mice were acclimatised for at least five days prior to experimental use.

*Eµ-Myc* transgenic mice (*30, 31*), always kept as heterozygotes, were maintained on a *C57BL/6- LY5.1-WEHI* background.

*dCas9a-SAM* transgenic mice (*8*) were maintained on a *C57BL/6-LY5.2-WEHI* background.

### Cell culture

*Eμ-Myc^T/+^;dCas9a-SAM^KI/+^:sgBcl2* cells (DHL214/DHL216) were derived from the haematopoietic organs of a wild-type mouse that previously underwent haematopoietic reconstitution with *Eμ- Myc^T/+^;dCas9a-SAM^KI/+^:sgBcl2* foetal liver cells (*8*). DHL214/DHL216 cells were maintained in cell culture using “FMA” media (500 mL high-glucose Dulbecco’s modified Eagle’s medium (DMEM), supplemented with 10% (50 mL) foetal bovine serum (FBS, Sigma-Aldrich #F9423), 50 μM 2- mercaptoethanol (Sigma-Aldrich #M3148) and 100 mM asparagine (Sigma-Aldrich #A4284), and were cultured for no longer than 3 months.

*Eμ-Myc^T/+^;dCas9a-SAM^KI/+^* cells (*Eμ-Myc312*) were derived from a transgenic *Eμ-Myc^T/+^;dCas9a- SAM^KI/+^* mouse that succumbed to lymphoma. *Eμ-Myc312* cells were maintained in cell culture under the same conditions as DHL214/DHL216 cells.

While being transduced with sgRNAs (individual or libraries), FLCs were maintained in foetal liver media: α-MEM with GlutaMax (Gibco #32561037), 10% FBS, HEPES (10 mM; Gibco #15630- 080), additional GlutaMax (1x; Gibco #35050-061) sodium pyruvate (1 mM; Gibco #11360-070), and β-mercaptoethanol (50 µM; Sigma-Aldrich #M3148), supplemented with mSCF (0.1 µg/mL; Peprotech #250-03), IL-6 (0.01 µg/mL; made in-house), TPO (0.05 µg/mL; Peprotech #315-14), and FLT-3 (0.01 µg/mL; made in-house).

HEK293T cells were maintained in cell culture using DMEM with 10% FBS and supplemented with penicillin (100U/mL) and streptomycin (100µg/mL) (Gibco #15140122).

Primary bone marrow derived macrophages (BMDMs) were first generated from *dCas9a- SAM^KI/KI^* mice as previously described (*23*). BMDMs were cultured in DMEM supplemented with 10% FBS, 15% L929-conditioned DMEM, 100 U/mL penicillin, and 100 mg/mL streptomycin for 3-4 days. BMDMs were immortalised by incubating the cells for 3-4 days in L929-conditioned medium containing *Cre-J2* retrovirus (from *Cre-J2* virus-producing cells (*23*)). Media was then removed and replaced with fresh L929-conditioned medium containing *Cre-J2* retrovirus, and cells incubated for a further 3-4 days. Media was then again removed and the cells cultured in complete L929-supplemented DMEM for 3-4 days until confluence. The percentage of L929- conditioned medium in the complete medium was slowly reduced over 2-3 months until cells could survive in the absence of L929-conditioned medium. These *dCas9a-SAM^KI/KI^* immortalised BMDMs (iBMDMs) were then maintained in DMEM supplemented with 10% FBS and 100 U/mL penicillin at 37°C and 5% CO_2_.

Cell lines were cultured at 37°C with 10% CO_2_ and regularly tested for Mycoplasma infection using a MycoALert detection kit (Lonza #LT07-118).

### Virus production and cell transduction

DHL214/DHL216 cells were derived from mice that underwent haematopoietic reconstitution with *Eµ-Myc^T/+^;dCas9a-SAM^KI/+^:sgBcl2* foetal liver cells, and were previously generated and characterised (*8*). Transduction of the 6 CRISPRa sgRNA libraries was undertaken by generating HEK293T cells (ATCC #CRL-3216) transfected with the library of choice (10 µg) and the lentiviral packaging constructs pMDL (5 μg), pRSV-REV (2.5 μg), and pVSV-G (3 μg) by calcium-phosphate precipitation, as previously described (*89, 90*). 48 h post-transfection, DHL214 cells were then transduced with supernatant containing viral particles (filtered using a 0.45 μm filter and supplemented with polybrene (8 μg/mL; made in-house)) via centrifugation (32°C, 2200 rpm, 2 h)).

For experiments involving iBMDMs, virus generation was undertaken as described above for DHL cells, and iBMDM transduction was performed by the direct addition of viral particles to culture dishes and incubation for 24 h with no centrifugation.

Foetal liver cells (FLCs) were transduced using plasmid DNA (10 μg; either sgRNA libraries 1-6 or individual sgRNAs) packaged with pMDL (5 μg), RSV-Rev (2.5 μg), and ENV (5 μg). Retronectin (32 µg/mL in 1× PBS)-coated tissue culture plates (Thermo Fisher Scientific #150200) were washed (1× PBS) and blocked (2% BSA in 1× PBS), and then the viral supernatant (48 h post-transfection) was filtered and applied to the plate by centrifugation. Supernatant was then removed and FLCs incubated in this plate overnight in foetal liver media (see above). Transduction efficiency for the FLCs was assessed via flow cytometry (based on BFP+ status) and found to be between ∼20-40%.

### Screening in iBMDMs

Partita sub-library 1-transduced iBMDMs were first treated with puromycin (8 µg/mL) for 4 days, to select for cells that were successfully transduced. After this selection, samples from each replicate were harvested for DNA (time point 0 (T0)). The puromycin-selected cells were then maintained in culture under normal conditions for 3 weeks. At the end of each week, cells from each replicate were harvested for DNA (time points 1/2/3 (T1/2/3)).

### CRISPRa enrichment screening in vitro in lymphoma cells with nutlin-3a, etoposide, and venetoclax

DHL214/216 cells were rapidly expanded for ∼10 days post-transduction, aiming to both avoid premature sgRNA selection while also allowing gene induction to begin. After ∼10 d, “input” cell pellets were collected, and DHL214/216 cells were then plated (in triplicate) into T75 flasks (libraries 1, 2, 3: ∼2x10^6^ cells, libraries 4, 5: ∼10x10^6^ cells). Each replicate was then treated with either DMSO (Sigma-Aldrich #D4540), nutlin-3a (Cayman Chemical #18585 or MedChemExpress #HY-10029), etoposide (Ebewe), or venetoclax (Active Biochem #A-1231). Treatment concentrations were sufficient to kill ≥90% of cells. Concentrations for screens using nutlin-3a and etoposide are shown in Table S1. Concentrations for screens using venetoclax differed between cell lines: DHL214 cells were treated with concentrations of 10 nM (IC_50_) and 30 nM (IC_80_), while DHL216 cells were treated with concentrations of 5 nM (IC_50_) and 15 nM (IC_80_). The nutlin-3a-treated samples for screens with sub-libraries 3, 4, and 5 were re-treated once post-initial recovery, using the same dose. Once the DHL214/216 cells had recovered to numbers judged sufficient for adequate DNA acquisition, cell pellets were collected.

### CRISPRa enrichment/depletion screening in vitro in iBMDMs

iBMDMs were transduced with Partita sub-library 1 lentiviral supernatant in the presence of 8 µg/mL polybrene. The infection was performed at a multiplicity of infection (MOI) < 0.5 to achieve 500x cell coverage per sgRNA, in triplicate. 500x cell coverage per sgRNA was maintained across the whole screening process. Library-transduced iBMDMs were selected with 8 µg/mL puromycin for 4 days. Time point 0 (T0) samples were collected following 4-day puromycin selection. Cells were then expanded and passaged every 2-3 days. Time points 1, 2, and 3 (T1, T2, T3) cell samples were collected at days 7, 14, and 21 post-puromycin selection, respectively.

### Foetal liver cell generation and transplantations for in vivo screening and validations

*dCas9a-SAM^KI/KI^* female mice were crossed with *Eµ-Myc^T/+^* male mice in timed matings to generate *dCas9a-SAM^KI/+^;Eµ-Myc^T/+^* E13.5-E14.5 embryos. The foetal livers were harvested from these embryos, and the foetal liver cells (FLCs) dissociated into single-cell suspensions via trituration, then stored at -80°C in a solution of 90% FBS and 10% DMSO (Sigma-Aldrich #D8418). Male *C57BL/6-LY5.1-WEHI* mice (6-8 weeks old) were lethally irradiated with two doses of 5.5 Gy γ-radiation (for reconstitution with *Eμ-Myc* transgene-containing foetal liver cells). Mice were randomly assigned to experimental groups, and then intravenously injected with ≥2×10^6^ FLCs (transduced with sgRNAs as described above) in 200 μL 1× PBS that had been washed and filtered. Mice were administered neomycin (2 mg/mL, Sigma-Aldrich #N1876) *ad libitum* via their drinking water for up to 4 weeks post-transplantation. Mice were monitored for lymphoma development by trained technicians blinded to the nature of the experiment. At ethical endpoint, peripheral blood was collected and analysed via Advia (Siemens), and tumour-burdened tissues were harvested, weighed, and both frozen as single cell suspensions and plated for cell line generation after homogenising tissues through a 100 µM strainer. Tumour-free survival curves, blood content, and organ weight data were statistically analysed using Prism (GraphPad). Tumour-free survival was calculated based on time between FLC transplantation and ethical endpoint. Mice were excluded from tumour-free survival analyses (and additional NGS/bioinformatics) if determined to have reached ethical endpoint due to unrelated reasons (e.g. skin infection).

### Next-generation sequencing and bioinformatic analyses

Genomic DNA (gDNA) was extracted from 4x10^6^ DHL cells or 5x10^6^ lymphoma-burdened splenocytes using the DNeasy Blood and Tissue kit (QIAGEN #69504) or Puregene Cell Kit (QIAGEN #158043) as per the manufacturer’s instructions. DNA content was quantified using a Nanodrop spectrophotometer (Denovix DS-11). All sgRNAs within the gDNA samples were then amplified and indexed in a single PCR, using 100illng of DNA and GoTaq Green Master Mix (Promega, #M7123). Primers were used which had been modified to possess unique 5’ overhangs/indexes compatible with Illumina sequencing (Table S6). The following PCR protocol was used: 3 mins at 95°C, [15 secs at 95°C, 30 secs at 60°C, 30 secs at 72°C repeated 35 times], and 7 mins at 72°C. For DHL screen samples, PCRs for each sample were performed in duplicate/triplicate, while *in vivo* sample PCRs were performed once. PCR products were pooled, cleaned up using Ampure XP beads (Beckman Coulter #A63881), and sequenced using a NextSeq 2000 (Illumina) as per the manufacturer’s instructions.

For iBMDMs, genomic DNA was isolated from samples using a Puregene Cell Kit (QIAGEN #158043). sgRNAs were amplified from 84 µg of DNA from each sample using Ultra II Q5 Master Mix (New England Biolabs #M0544) according to the manufacturer’s instructions. Primers used for sgRNA amplification are shown in Table S6. 3 unique library indexes and 7 reactions were used to amplify each genomic DNA sample. Amplified and indexed PCR products were pooled and purified using 0.9 volumes of AMPure XP Beads (Beckman Coulter #A63880). The sequencing was performed using NextSeq 2000 P3 XLEAP-SBS Reagents (100 cycles) on the NextSeq 2000 system (Illumina) according to the manufacturer’s instructions.

### Statistical analyses for the double-hit lymphoma and in vivo enrichment screens

For the screens in DHL cells using nutlin-3a/etoposide, MAGeCK v0.5 (37) was used to rank and compare sgRNA enrichment within the different treatment samples.

For the in vivo screening, MAGeCK v0.5 was used to determine the representation of sgRNAs per tumour, as calculated by the number of reads mapping to each sgRNA in the library within a sample.

Raw sequencing data is available on the NCBI GEO repository (GSE296642).

### Statistical analyses for the iBMDM enrichment/depletion screen

Reads were mapped to the libraries and raw count data generated using MAGeCK v0.5.9 (*37*). Raw count data were then stored in a standard Bioconductor SummarizedExperiment object (*91*) and normalized for sequencing depth. We performed a differential abundance analysis for each sgRNA separately using the limma-voom approach (*92*). Specifically, we fit a linear model to the log-CPM values for each sgRNA, using voom-derived observation and quality weights. We performed robust empirical Bayes shrinkage to obtain shrunken variance estimates for each sgRNA, and we used moderated F-tests to compute p-values for each of the two-group comparisons of interest. To obtain promoter-level statistics, we used the “fry” gene-set enrichment analysis (GSEA) method implemented in limma, and considered all sgRNAs targeting a given promoter as a “gene set”. This allowed for the detection of enriched or depleted promoters. We then applied the Benjamini-Hochberg procedure to obtain an FDR-corrected p-value for each promoter. Essential/non-essential genes were determined as previously described (*93*).

To perform GSEAs, a hypergeometric test was used separately for enriched genes (logFC>2) and depleted genes (logFC<-2) at each time point, using the initial time point as the reference control group. Gene ontology (GO) data for murine gene sets were obtained from the MSigDB database (*94*).

Raw sequencing data is available on the NCBI GEO repository (GSE296642).

### Individual sgRNA cloning and transduction

Custom sgRNAs were ordered from Integrated DNA Technologies (sequences in Table S4), with the sequence taken from the most highly enriched sgRNA for that gene in the screens. The sgRNAs were modified to possess overhang sequences (sense: 5’-ACCG-3’, antisense: 5’-AAAC- 3’), facilitating cloning into the BsmBI site of our constitutive expression WEHI-12 vector. A non-targeting control sgRNA (*sgNTC*) was also generated (Table S4). Successful sgRNA insertion into WEHI-12 was confirmed via sequencing (service provided by the Australian Genome Research Facility), using the primer 5’- GAGGGCCTATTTCCCATGATT-3’. For the *in vivo* screening and validations, a positive control guide targeting *Bcl2* (*sgBcl2*) was used (Table S4), which was cloned into LV06 (Sigma-Aldrich). Once cloned, each sgRNA vector was purified via phenol/chloroform extraction, transformed into electrocompetent Stbl4 cells (Thermo Fisher #11635018). The DNA was then extracted, using a either a PureLink HiPure Plasmid DNA Purification kit (Invitrogen #K210005) or a Plasmid Purification Kit (QIAGEN #12145), and transduced into DHL214 cells (as described above). Stably transduced DHL214 cells were then sorted out using a FACSAria III or FACSAria Fusion (BD Biosciences).

### Cell viability assays for individual sgRNA validation

DHL214 cells were seeded onto 96 well plates at 40,000 cells per well, in technical duplicate, then treated with either DMSO, nutlin-3a (MedChemExpress #HY-10029), etoposide (Ebewe #EB461), A-1331852 (a gift from Dr N Duong and Prof G Lessene (WEHI, Australia)), S63845 (Chemgood #C1370), or venetoclax (Active Biochem #A-1231) at the indicated concentrations for 24 h. Cells were washed in PBS and resuspended in 1x Annexin V binding buffer (10x recipe: 0.1 M pH 7.4 HEPES, 1.4 M NaCl, 25 mM CaCl_2_, made in PBS) containing PI (1 μg/mL, Sigma-Aldrich #P4170) and Annexin V-A647 (1:1000-2000, made in house). Cells were analysed via FACS using an LSR IIW (BD Biosciences), and data were analysed using FlowJo (BD Biosciences) and Prism 9 (GraphPad). Each viability assay was performed at least three times.

### Competition assays for individual sgRNA validation

DHL214 cells transduced with *sgNTC* were further transduced (as above) with FUGW (Addgene #14883), as the FUGW vector constitutively expresses GFP (more strongly than the endogenous *dCas9-SAM* alone) which makes the eventual *sgNTC*+FUGW cells easily distinguishable. *sgNTC*+FUGW DHL214 cells were then compared to each other cell line by seeding them each in 6 well plates at 200,000 cells per well, per cell line, with technical duplicates. Cells were treated at approximate IC_20_ concentrations (as determined by the *sgNTC* DHL214 cells) with either DMSO, nutlin-3A (1.2 μM) or etoposide (7 ng/mL), and then passaged every 2-3 days, with fresh drug applied at each passage. The proportions of *sgNTC*+FUGW cells and experimental cells were quantified via FACS using an LSRFortessa X-20 (BD Biosciences) at day 0, and then intervals of 2-3 days over 14 days. An additional biological replicate was performed for the control assay (*sgNTC*+FUGW vs. *sgNTC*) and for each sgRNA considered to potentially have validated after the first experiment. Data were analysed using FlowJo (BD Biosciences) and Prism 9 (GraphPad).

### Quantitative reverse transcriptase PCR

RNA was extracted from cells frozen in TRIzol (Thermo Fisher Scientific #15596018) as per manufacturer’s instructions. cDNA was synthesised using a SuperScript III First Strand Synthesis system (Thermo Fisher Scientific #11904018 or #11752050) as per manufacturer’s instructions. qRT-PCR assays were performed using TaqMan Fast Advanced Master Mix (Thermo Fisher Scientific #4444557), and TaqMan Assay probes targeting various genes (Table S5, all from Thermo Fisher Scientific), each according to the manufacturer’s instructions. All qRT-PCRs used 3 technical replicates per sample, with biological replicates indicated in their respective figure legend. All qRT-PCRs were run on a QuantStudio 12K Flex Real-Time PCR System (Thermo Fisher Scientific). Data were analysed using the ΔΔCt method, with each sample normalised to expression of a housekeeping control gene (*Hmbs*) and their respective control samples (indicated in the figure legend for each experiment).

### Statistical analyses

All graphs and statistical analyses were generated and performed using Prism software (v8+; GraphPad). All statistical tests used are indicated in the legend of the relevant figure. Data are presented as mean ± standard deviation (SD), unless otherwise stated. Statistically significant differences between groups was ascribed when a p-value was less than 0.05.

### Flow cytometry and immunophenotyping

After euthanasia of reconstituted mice, bone marrow was flushed from femurs and tibias with PBS containing 5% FBS and 5 µM EDTA using a 1 mL syringe and 25 G needle, and then gently dissociated into a single cell suspension with a P1000 pipette. Single cell suspensions were cryopreserved in FCS containing 10% DMSO. For immunophenotyping by flow cytometry, bone marrow cells were thawed and ∼5x10^6^ cells were live/dead stained with ViaDye Red (Cytek #R7- 60008) according to manufacturer’s instructions. Cells were then stained for 1 h on ice with a cocktail of fluorochrome conjugated antibodies (Table S6) diluted in PBS containing 5% FCS and 10% FcG receptor antibody-containing supernatant (made in-house; for blocking non-specific binding). Stained cells were analysed on an Aurora cytometer (Cytek). Data was analysed using FlowJo software (BD Biosciences). Doublets were excluded by FSC-A/FSC-H, live cells were gated on by ViaDye Red exclusion and GFP+, debris was excluded by gating on FSC-A/SSC-A, lymphocytes containing sgRNA were selected by GFP+/BFP+ fluorescence. B cell immunophenotype was determined by B220 and CD19 expression, and then by surface IgM and/or IgD expression.

### Western blotting

Cell pellets were lysed in radio-immunoprecipitation assay (RIPA) buffer in the presence of cOmplete protease inhibitors (Merck #311697498001), and a Pierce BCA Protein assay (Thermo Fisher Scientific #23225) was performed to quantify protein. 10 µg of protein was loaded onto 4-12% Bis-Tris NuPAGE gels (Thermo Fisher Scientific #NP0322BOX). Protein markers used were either a Spectra Multicolor Broad Range Protein Ladder (Thermo Fisher #26634) or a Precision Plus Protein Kaleidoscope Prestained Protein Standard (Bio-Rad #1610375). Proteins were transferred to PVDF membranes (Thermo Fisher Scientific #IB24001) using the iBlot2 system (BioRad). Membranes were blocked in 5% skim milk in TRIS-buffered phosphate containing 1% TWEEN-20 (TBST) for 1 h at room temperature. Membranes were washed in TBST and incubated in primary antibodies diluted in 5% bovine serum albumin (BSA) in TBST overnight at 4°C. Primary antibodies used are: mouse anti-HSP70 (1:10,000, gift from Prof R Anderson (ONJCRI, Australia), clone: N6), rabbit anti-BIM (1:1000, Enzo Life Sciences #ADI-AAP-330-E, RRID: AB_2038875, clone: polyclonal), mouse anti-BCL-2 (1:2000, BD Biosciences #610539, RRID: AB_397896, clone: 7/BCL-2), rabbit anti-PUMA (1:500, ProSci Incorporated #3043, RRID: AB_203251, clone: polyclonal), rabbit anti-TRP53 (1:2000, Novocastra #NCL-p53-CM5p, RRID: AB_563933, clone: CM5), rabbit anti-P19ARF (1:1000, Sigma-Aldrich #PC435, RRID: AB_213725), rat anti-BCL-XL (1:1000, made in-house, clone: 9C9), rabbit anti-A1 (1:500, made in-house, clone: 6D6), and rabbit anti-MCL-1 (1:1000, made in-house, clone: 14C11-20). Membranes were then washed and incubated in secondary antibodies diluted in 5% skim milk in TBST for 1 h at room temperature. Secondary antibodies used are: goat anti-mouse IgG (1:2000, Southern Biotech #1010-05, RRID: AB_2728714), goat anti-rabbit IgG (1.25:5000, Southern Biotech #4010-05, RRID: AB_2632593), and goat anti-mouse IgG (1:5000, Southern Biotech #3010-05, RRID: AB_2795801). Finally, membranes were washed and imaged with Immuno Forte Chemiluminescent reagent on a Chemidoc (BioRad).

## Supplementary Materials and Tables

Supplementary File 1. Complete information for Partita sub-libraries 1-6, including sgRNA ID numbers, sgRNA sequences, and the gene targeted by each sgRNA.

Supplementary File 2. Uncropped western blots for those included in this study.

**Supplementary Table 1.** Concentrations of nutlin-3a and etoposide used to treat DHL214 cells in the *in vitro* screens.

| Partita sub-libraries | Treatments | Concentrations |
| --- | --- | --- |
| sub-library 1 | nutlin-3A | 1 μM |
|  | etoposide | 20 ng/mL |
| sub-library 2 | nutlin-3A | 1 μM |
|  | etoposide | 20 ng/mL |
| sub-library 3 | nutlin-3A | 1 μM |
|  | etoposide | 10 ng/mL |
| sub-library 4 | nutlin-3A | 2 μM |
|  | etoposide | 10 ng/mL |
| sub-library 5 | nutlin-3A | 2 μM |
|  | etoposide | 10 ng/mL |

**Supplementary Table 2.** Hit genes from *in vitro* screens in DHL cells using venetoclax, chosen for independent validation experiments.

| Gene (mouse) | Full gene name (mouse) | Sub-library of origin | Validated after 24 h viability assay (as assessed by IC<sub>50</sub> fold change)? |
| --- | --- | --- | --- |
| *Dyrk1a* | *dual specificity tyrosine phosphorylation regulated kinase 1A* | 1 | N |
| *Irx5* | *iroquois homeobox 5* | 2 | Y |
| *Phgdh* | *3-phosphoglycerate dehydrogenase* | 2 | N |
| *Rac1* | *Rac family small GTPase 1* | 2 | Y |
| *Trip6* | *thyroid hormone receptor interactor 6* | 2 | Y |

**Supplementary Table 3.** Hit genes from *in vitro* screens in DHL cells using nutlin-3a and etoposide, chosen for independent validation experiments.

| Gene (mouse) | Full gene name (mouse) | Sub-library of origin | Validated after 24 h viability assay? | Validated after 14 d competition assay? |
| --- | --- | --- | --- | --- |
| *Rnf128* | *ring finger protein 128* | 1 | N | N |
| *Hira* | *histone cell cycle regulator* | 1 | Y | Y |
| *Aoc2* | *amine oxidase copper containing 2* | 1 | N | Y |
| *Pik3c2a* | *phosphatidylinositol-4-phosphate 3-kinase catalytic subunit type 2 alpha* | 1 | Y | Y |
| *Hic2* | *hypermethylated in cancer 2* | 1 | N | N |
| *Ssxb10* | *SSX member B10* | 2 | N | N |
| *Phgdh* | *3-phosphoglycerate dehydrogenase* | 2 | Y | Y |
| *Dnaja1* | *DnaJ heat shock protein family (Hsp40) member A1* | 2 | Y | Y |
| *Gsdma* | *gasdermin A* | 3 | N | N |

**Supplementary Table 4.**
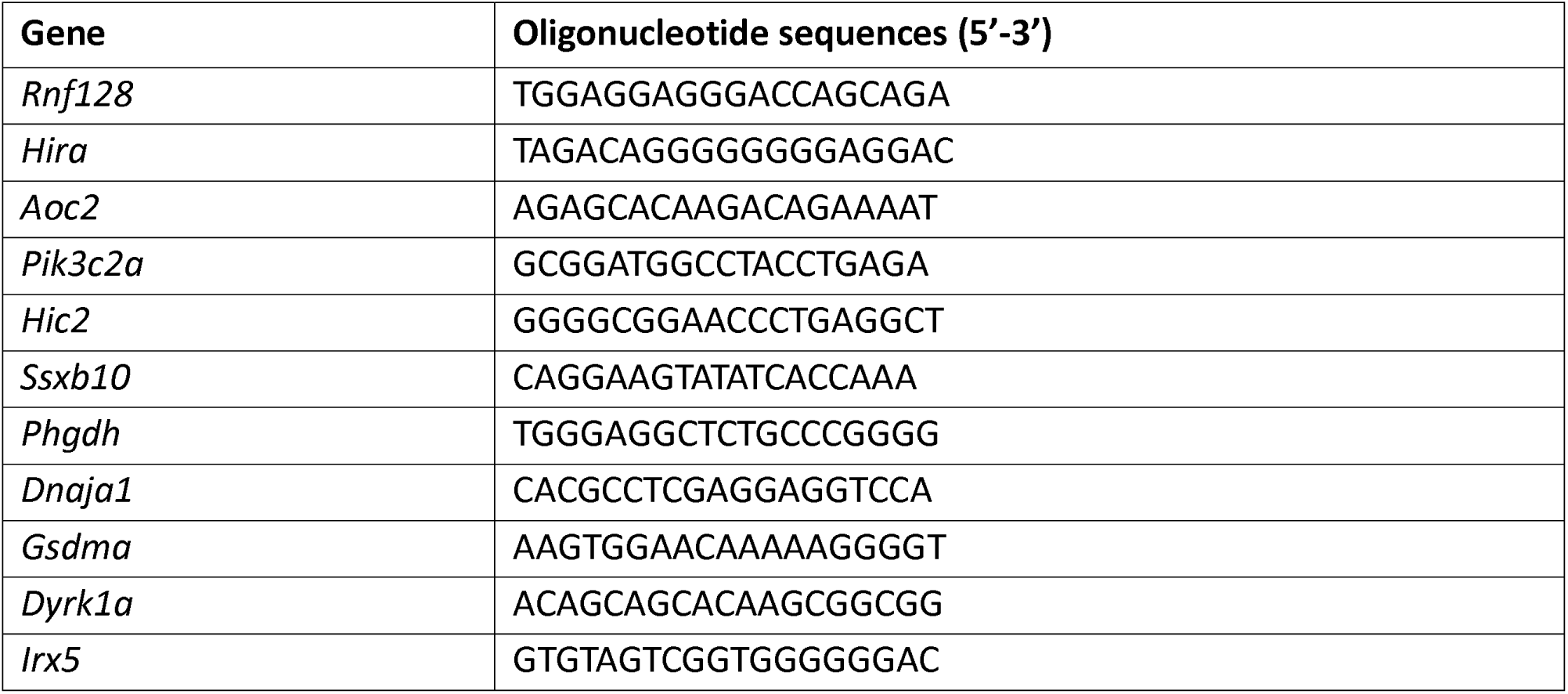
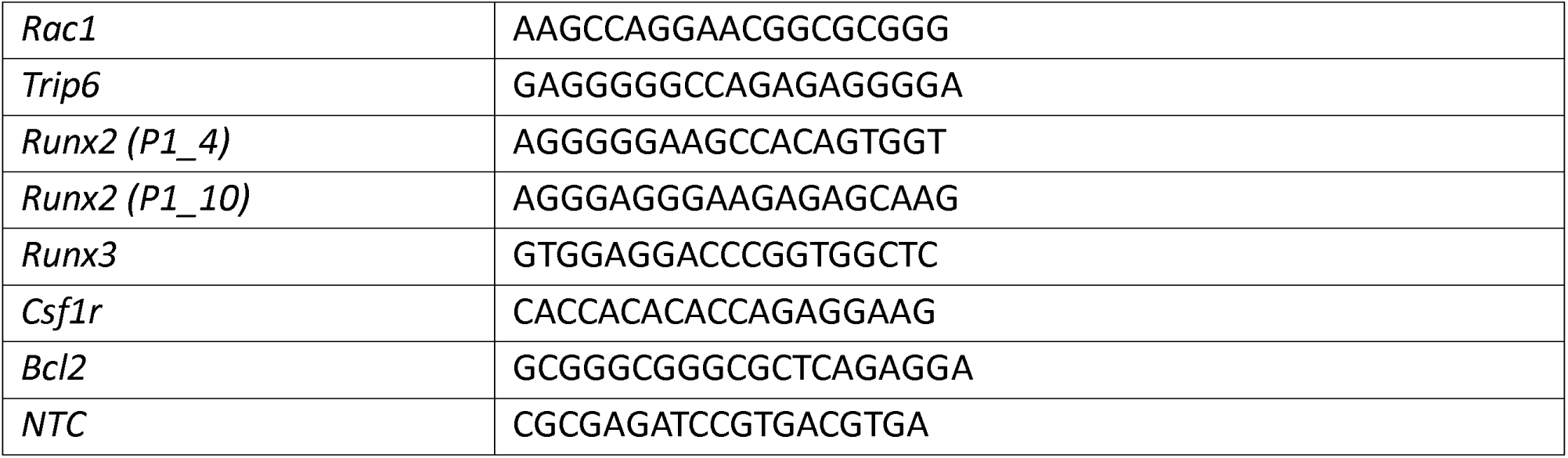
Oligonucleotide sequences used to generate individual sgRNAs for gene validation experiments. Overhangs to clone the oligos into the BsmBI site of the WEHI-12 vector were included for each (sense: 5’-ACCG-3’, antisense: 5’-AAAC-3’). The sequences of the NTC and Bcl2 sgRNAs used (in all relevant experiments) is also shown. All sequences are taken from the Partita library.

**Supplementary Table 5.** Identification numbers of the TaqMan probes used to assess gene upregulation during validations via qRT-PCR.

| Target Gene | TaqMan probe ID |
| --- | --- |
| *Rnf128* | Mm00480990_m1 |
| *Hira* | Mm00468902_m1 |
| *Aoc2* | Mm00841716_m1 |
| *Pik3c2a* | Mm00478162_m1 |
| *Hic2* | Mm05914754_s1 |
| *Ssxb10* (has off-targets in *Ssxb2* and *Ssxb8*) | Mm04212947_gH |
| *Phgdh* | Mm01623589_g1 |
| *Dnaja1* | Mm00787254_s1 |
| *Gsdma* | Mm01306778_m1 |
| *Hmbs* | Mm01143545_m1 |
| *Bcl2* | Mm00477631_m1 |
| *Csf1r* | Mm01266652_m1 |
| *Runx2* | Mm005011584_m1 |
| *Runx3* | Mm00490666_m1 |
| *Irx5* | Mm00502107_m1 |
| *Rac1* | Mm01201653_mH |
| *Trip6* | Mm00600041_m1 |

**Supplementary Table 6.**
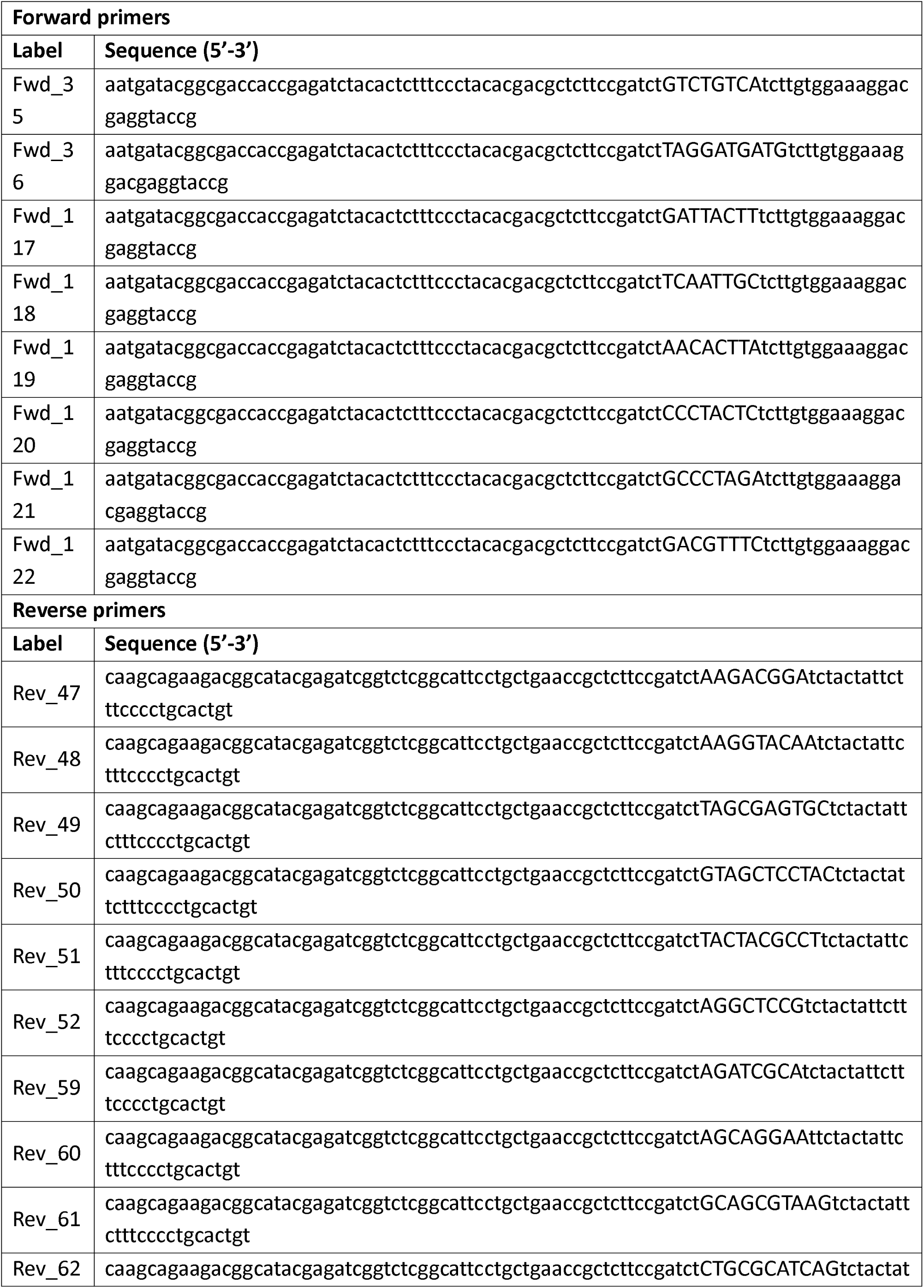
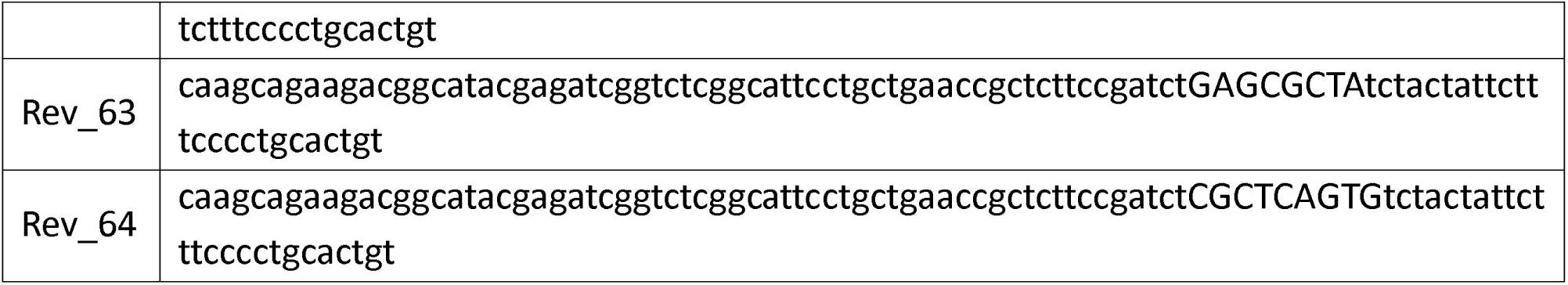
Overhang/indexing primers used in this study for NGS of the Partita library. The barcode sequence is shown in upper case, while the Illumina overhang sequence is in lowercase at the 5’ end, and the vector-binding sequence is in lowercase at the 3’ end. The primers are compatible in any combination of forward/reverse, and were used as such, while ensuring that no two samples were amplified using the same primer combination.

**Supplementary Table 6.** Antibodies used for flow cytometric immunophenotypic analyses. Abbreviations: APC = allophycocyanin; BV = Brilliant Violet; PE = phycoerythrin.

| Antigen | Clone | Fluorescent conjugate | Supplier | Catalogue number |
| --- | --- | --- | --- | --- |
| B220 | RA3-6B2 | BV605 | BioLegend | #103244 |
| IgM | 5.1 | PE | made in-house | N/A |
| IgD | 11-26C.2a | BV510 | BD Biosciences | #563110 |
| CD19 | ID3 | APC | made in-house | N/A |
| ViaDye Red | N/A | N/A | Cytek | #SKU R7-60008 |

## Acknowledgements

STD performed and bioinformatically analysed the *in vitro* screens with DHLs using venetoclax. All associated validations of these screens were performed by STD, with assistance from CC and JELM. YD performed the *in vitro* screen with iBMDMs. Bioinformatic analyses of the iBMDM screen were performed by JPF. MAP performed the *in vivo* screen, and performed the bioinformatic analyses with assistance from STD. Validations of individual genes *in vivo* were undertaken by JELM, with assistance from STD, CK, and MAP. JELM performed the *in vitro* screens with DHLs using nutlin-3a and etoposide, which were bioinformatically analysed by JELM and STD. All associated validations of these screens were performed by JELM, with assistance from CC, FB, and STD.AH, KD, JPF, and BH designed and generated the Partita library. AH led the process of sgRNA targeting and overall library optimisation. KD designed the D5 scaffold and contributed to vector optimisation. AK assisted with the design and optimisation of PCR and NGS protocols. LW performed all live mouse-associated techniques. LT performed and/or assisted with all cloning techniques. MJH designed and supervised the study with input from BH and GLK. JELM drafted manuscript with assistance from MAP, YD, and STD. JELM revised the manuscript. JELM prepared the figures, with assistance from MAP, YD, and STD. All authors reviewed and approved the manuscript.

We thank Marcel Doerflinger for advice relating to BMDM immortalisation. We thank Colin Watanabe for assistance with the initial library designs and TSS mapping. We thank Andreas Strasser for helpful comments on the manuscript. We thank the WEHI BioServices facility for assistance with all *in vivo* experiments. We thank Stephen Wilcox, Sarah MacRaild, and the WEHI genomics facility for assistance with all NGS experiments and data management. We thank the FACS facilities at both the ONJCRI and WEHI.

This work was supported by grants/awards to: STD (Victorian Cancer Agency grant 21006), JELM (Cancer Council Victoria Grant-in-Aid, Robert and Janette Boffey donation), GLK (NHMRC grants GNT2002618, GNT2001201, and GNT2011139 (with Andrew Roberts and Andrew Wei); Victorian Cancer Agency grant 17028; donation from the estate of Anthony (Toni) Redstone OAM (with Andreas Strasser); a donation from the Craig Perkins Cancer Research Foundation; a donation from the Dyson Bequest; and a donation from the Harry Secomb Foundation), MJH (The Australian Lions Childhood Cancer Research Foundation Grant (with Emily Lelliott); NHMRC grants 2017971, GNT1159658, GNT1186575, GNT1145728, and GNT1143105; and a Cancer Council Victoria Venture Grant).

JPF and BH are former employees of Genentech, and AH and KD are current employees of Genentech, where the Partita library presented in this work was developed. The remaining authors declare that they have no competing interests.

All data needed to evaluate the conclusions in the paper are present in the paper and/or the Supplementary Materials. Additional raw sequencing data is available on the NCBI GEO repository (GSE296642). Data or materials can be provided by the authors pending scientific review and, where necessary, a completed material transfer agreement. Requests for the data/materials should be submitted to the corresponding author(s).

